# Insoluble Aβ overexpression in an *App* knock-in mouse model alters microstructure and gamma oscillations in the prefrontal cortex, causing anxiety-related behaviours

**DOI:** 10.1101/494443

**Authors:** Eleftheria Pervolaraki, Stephen P Hall, Denise Foresteire, Takashi Saito, Takaomi C Saido, Miles A Whittington, Colin Lever, James Dachtler

## Abstract

We studied two new *App* knock-in mice models of Alzheimer’s disease (*App^NL-F^* and *App^NL-G-F^*), which generate elevated levels of Aβ_40_ and Aβ_42_ without the confounds associated with APP overexpression. This enabled us to assess changes in anxiety-related and social behaviours, and neural alterations potentially underlying such changes, driven specifically by Aβ accumulation. *App^NL-G-F^* knock-in mice exhibited subtle deficits in tasks assessing social olfaction, but not in social motivation tasks. In anxiety-assessing tasks, *App^NL-G-F^* knock-in mice exhibited: 1) increased thigmotaxis in the Open Field (OF), yet; 2) reduced closed-arm, and increased open-arm, time in the Elevated Plus Maze (EPM). Their ostensibly-anxiogenic OF profile, yet ostensibly-anxiolytic EPM profile, could hint at altered cortical mechanisms affecting decision-making (e.g. ‘disinhibition’), rather than simple core deficits in emotional motivation. Consistent with this possibility, alterations in microstructure, glutamatergic-dependent gamma oscillations, and glutamatergic gene expression were all observed in the prefrontal cortex, but not the amygdala, of *App^NL-G-F^* knock-in mice. Thus, insoluble Aβ overexpression drives prefrontal cortical alterations, potentially underlying changes in social and anxiety-related behavioural tasks.

## Introduction

Alzheimer’s disease (AD) is classically associated with declining cognitive function (Scheltens et al., 2016). However, this is only one aspect of the behavioural changes associated with AD. Other behavioural changes include reduced social engagement and increased anxiety. Although social aspects of AD have remained underexplored, social withdrawal is present up to 5 years prior to a clinical cognitive diagnosis (Jost and Grossberg, 1995). AD patients with larger social networks (the number of people with which one has meaningful contact with) have slower cognitive decline, compared to AD patients with small social networks (Bennett et al., 2006). Social factors may modulate the rate of disease pathology and cognitive decline, but crucially, also the chance of developing AD (Kuiper et al., 2015). Studies have found that elderly people who identify as lonely, the risk of developing AD was nearly doubled (Wilson et al., 2007). Recent evidence from the Lancet Commission Report highlights that social isolation constitutes a 2.3% risk of developing AD (Livingston et al., 2017). Thus, social factors not only modulate the risk of developing dementia, but also disease progression. Together, it can be inferred that changes in social motivation of the individual, as a result of AD pathology, may be a factor in disease progression.

Anxiety in AD is relatively common, with up to 71% of patients reporting anxiety concerns (Ferretti et al., 2001; Teri et al., 1999). Up to 6% had anxiety that reached the diagnostic criteria of generalised anxiety disorder of Diagnostic and Statistical Manual of Mental Disorders (Ferretti et al., 2001). Anxiety behaviours may also predict conversion to AD. Mild cognitive impairment (MCI) is often the precursor condition to AD. 83.3% of MCI patients that also exhibited anxiety symptoms converted to AD within a three year follow up period compared to 40.9% of persons with MCI but without anxiety (Palmer et al., 2007). This and other studies (Gallagher et al., 2011; Li and Li, 2018; Pietrzak et al., 2015) suggest that anxiety is associated with early phases of AD. Neurodegeneration in early AD may explain the increase in anxiety. The presence of anxiety is associated with abnormal CSF Aβ_42_ and tau in MCI patients (Ramakers et al., 2013), and thus may reflect underlying pathology. In summary, social-related and anxiety-related changes in AD can occur early in, and predict, disease progression.

The AD-affected neural regions relevant to changes in anxiety and social behaviours remain unclear but candidate regions of interest (ROIs) for the present study were the hippocampus, amygdala, and prefrontal cortex. Our rationale for these ROIs was twofold. First, these regions have been consistently linked to anxiety and social behaviours (e.g., hippocampus: (Bannerman et al., 2004), (Hitti and Siegelbaum, 2014; Okuyama et al., 2016); amygdala (Shin and Liberzon, 2010); prefrontal cortex; (Cao et al., 2018)). Second, pathology in these regions can occur early in AD and predict MCI-to-AD conversion (e.g., hippocampus: (Devanand et al., 2007; Liu et al., 2010); amygdala (Liu et al., 2010; Poulin et al., 2011); prefrontal cortex: (Okello et al., 2009; Plant et al., 2010; Tondelli et al., 2012)). Interestingly, in Okello et al. (2009), anterior cingulate had the highest amyloid-beta load of six ROIs in MCI patients relative to controls, and anterior cingulate amyloid-beta load predicted faster conversion from MCI to AD.

Currently, we have few insights into the biological mechanisms underpinning anxiety and social withdrawal in AD, highlighting the need to find AD mice that model these behaviours.

Broadly speaking, pathologies underpinning AD tend to group into two domains; those of amylogenic pathways, and those of tau; although the two are not mutually exclusively (Bloom, 2014). The animal models have tended to focus on only one of these pathology pathways, although the majority of these model the amyloid aspects of AD through amyloid-β peptide (Aβ) plaques. Despite many transgenic mouse models of AD existing, previous generations of Aβ mice have achieved their Aβ overexpression by also overexpressing amyloid precursor protein (APP). Over time, it has become clear the APP overexpression alone can introduce confounds that make it difficult to dissociate the causal effects of Aβ compared to APP. Mice overexpressing human wild-type APP have cognitive impairments in the Morris watermaze and object recognition test, increased tau hyperphosphorylation and reduced GluA1, GluA2, GluN1, GluN2A, GluN2B, phosphorylated CaMKII and PSD95 (Simon et al., 2009). This was also associated with reduced cell density in the pyramidal layer of CA1 (Simon et al., 2009). Others have found that other Aβ mice (the APP23) produced higher amounts of APP fragments, including C-terminal fragment β/α and APP intracellular domain (Saito et al., 2014). The phenotypes associated with APP overexpression alone encapsulate what are known to be AD-specific pathologies, underlining the potential risks of using APP overexpression models to make AD interpretations.

Recently, a new generation of AD mouse models have become available that achieves Aβ pathologies without the overexpression of APP (Saito et al., 2014). This has been achieved by humanising the murine Aβ sequence through a knock-in (KI) strategy. Up to three familial mutations have been knocked-in to generate two AD mouse models. The *App^NL-F^* mouse contains the Swedish KM670/671NL mutation and the Iberian I716F mutation which results in elevated total Aβ and elevated Aβ_42_/Aβ_40_ ratio (Saito et al., 2014). The *App^NL-G-F^* mouse also contains the Swedish and Iberian mutations, but with the additional Artic mutation E693G which promotes aggressive Aβ oligomerisation and amyloidosis (Saito et al., 2014). In *App^NL-G-F^* mice, Aβ plaques are saturated by 9 months of age, but by contrast, *App^NL-F^* mice exhibit relatively few plaques. Both models have approximately similar amounts of soluble Aβ species (Saito et al., 2014). Hence, these two mouse models are useful to be able to dissociate the effects of soluble Aβ and plaque-based Aβ in the generation of AD-related pathologies.

Within the current study, we have sought to explore whether these new *App* KI mouse models display alterations in behaviours related to social motivation, social memory and anxiety. Briefly, in terms of behaviour, we found marked, yet seemingly contradictory, changes in anxiety tasks (‘anxiogenic’ in the OF, ‘anxiolytic’ in the EPM), together with mild deficits on social tasks, in *App^NL-G-F^* knock-in mice. Arguably, the marked anxiety-related behavioural changes pointed more towards changes in decision-making than towards a simple shift in core anxiety-related emotionality. Accordingly, we applied three methodological assays to these mice, probing aspects of anatomy, physiology, and genetics, with each assay consistently targeting both the prefrontal cortex (long linked to decision-making interfacing affect and cognition) and the basolateral amygdala (long linked to anxiety-related emotionality). Consistent with decision-making being affected more than core emotionality, (albeit with other interpretations possible), we found that the prefrontal cortex exhibited alterations in all three of our assays (microstructure, glutamatergic-dependent gamma oscillations, and glutamatergic gene expression) while the basolateral amygdala exhibited alterations in none of them. We conclude that insoluble Aβ overexpression leads to alterations in prefrontal cortices (as well as the hippocampus), which could at least partially underlie changes in social and anxiety-related tasks.

## Materials and Methods

### Ethics

All procedures were approved by the Durham University and University of York Animal Ethical and Welfare Review Boards and were performed under UK Home Office Project and Personal Licenses in accordance with the Animals (Scientific Procedures) Act 1986.

### Mice

Full details of the animals can be found elsewhere (Saito et al., 2014). Upon arrival at Durham, mice were backcrossed once to the C57BL/6J line, which was the same background strain as the previous institution where they were bred. Homozygote *App^NL-F^* and *App^NL-G-F^* and wild-type littermate mice were bred in house by heterozygote pairings and were weaned at P21 and ear biopsied for genotyping. Genotyping protocols can be found elsewhere (Saito et al., 2014). All mice were housed with littermates.

### Experimental animals

#### Behaviour

21 wild-type (9 male (8.19 months ± 0.12), 11 female (8.10 months ± 0.04)), 19 *App^NL-F^* homozygotes (10 male (7.83 months ± 0.15), 9 female (7.64 months ± 0.14)) and 19 *App^NL-G-F^* homozygotes (10 male (7.93 months ± 0.17), 9 female (7.44 months ± 0.43)) were used in behavioural testing. Mice were transferred to the testing room and allowed at least 30 mins habituation prior to testing. All mice were transferred to the apparatus via cardboard tubes.

Testing procedures can be found elsewhere (Dachtler et al., 2014). All trials were recorded by Any-maze (Stoelting, Dublin, Ireland) tracking software. Mice undertook the following test in this order: open field, elevated plus maze, social approach, social recognition and social olfactory discrimination. In brief, mice were placed into a 44 cm^2^ white Perspex arena and allowed free ambulation for 30 mins. The floor of the arena was divided by Any-maze into an outer zone of 8 cm and a centre zone of 17.5 cm^2^. Time spent and entries into these zones, along with distance travelled were measured. The elevated plus maze and social approach, social recognition, and social olfaction discrimination was run as in (Dachtler et al., 2014), except that novel conspecifics and soiled bedding was sex matched with adult mice to the test subject. At the conclusion of testing, mice were either used for electrophysiology, killed by perfusion fixation with 4% paraformaldehyde in phosphate buffered saline (PBS) or killed by cervical dislocation for molecular biology.

### Diffusion Tensor MRI

8 wild-type (4 male (9.02 months ± 0.63), 4 female (9.43 months ± 0.30)) and 8 *App^NL-G-F^* homozygotes (4 male (8.88 months ± 0.22), 4 female (9.89 months ± 0.21)) were used in for MR imaging. Image acquisition has been described elsewhere (Pervolaraki et al., 2017; Pervolaraki et al., 2019). MR imaging was performed on a vertical 9.4 Tesla spectrometer (Bruker AVANCE II NMR, Ettlingen, Germany). During imaging, the samples were placed in custom-built MR-compatible tubes containing Fomblin Y (Sigma, Poole, Dorset, UK). The following acquisition protocol was used: TE: 35 ms, TR: 700 ms, 1 signal average. The field of view was set at 168 × 128 × 96, with a cubic resolution of 62.5 μm/pixel and a B value of 1625 s/mm with 30 directions.

The *ex vivo* mouse brain 3D diffusion-weighted images were reconstructed from the Bruker binary file using DSI Studio (http://dsi-studio.labsolver.org). Direction Encoded Colour Map (DEC) images were generated by combining the information from the primary eigenvectors, diffusion images and the fractional anisotropy (FA). Regions were extracted by manually segmenting orbitofrontal cortex (OFC), anterior cingulate cortex (ACC), hippocampal and whole amygdalar regions using a mouse brain atlas (Franklin and Paxinos, 2008). We chose to focus on the aforementioned brain regions as these have been linked to social behaviour (Pervolaraki et al., 2019). Additionally, the hippocampus (Bannerman et al., 2004; McHugh et al., 2004), the amygdala (Davis, 1992; Davis et al., 1994) and prefrontal cortical regions (including the ACC) (Davidson, 2002; Etkin et al., 2011) have all been linked to anxiety. Extraction of FA, mean diffusivity (MD), axial diffusivity (AD) and radial diffusivity (RD) was performed within selected segmented brain regions, with 3 100 μm sections (one anterior and one posterior to the segmented section) extracted. Full atlas-based description of the segmentation can be found in (Pervolaraki et al., 2019).

### Electrophysiology

Anterior cingulate or basolateral amygdala coronal slices (450 μm thick) were prepared from age-matched, 12 month old male wild-type (n=5) mice and *App^NL-G-F^* KI (n=5) mice. Animals were anesthetised with inhaled isoflurane, immediately followed by an intramuscular injection of ketamine (100 mg/kg) and xylazine (10 mg/kg). Animals were perfused intracardially with 50-100ml of modified artificial cerebrospinal fluid (ACSF), which was composed of the following (in mM): 252 sucrose, 3 KCl, 1.25 NaH_2_PO_4_, 24 NaHCO_3_, 2 MgSO_4_, 2 CaCl_2_, and 10 glucose. The brain was removed and submerged in cold (4–5°C) ACSF during dissection. Once dissected, slices were maintained at 32°C in a standard interface recording chamber containing oxygenated ACSF consisting of (in mM): 126 NaCl, 3 KCl, 1.25 NaH_2_PO_4_, 1 MgSO_4_, 1.2 CaCl_2_, 24 NaHCO_3_ and 10 glucose. Persistent, spontaneous gamma oscillations were generated through the bath application of 400 nM kainic acid (KA). Further perfusion, through bath application of 20 mM 3-(2-Carboxypiperazin-4-yl)propyl-1-phosphonic acid (CPP), was conducted to antagonise NMDA receptors. All salts were obtained from BDH Chemicals (Poole, UK) or Sigma-Aldrich (Poole, UK) and KA and CPP were obtained from BioTechne (Minnesota, USA).

Extracellular field recordings were obtained using borosilicate micropipettes (Harvard Apparatus) filled with ACSF and had resistances of 2-5 (MΩ). Recordings were bandpass filtered at 0.1Hz to 300Hz. Power spectra were derived from Fourier transform analysis of 60 s epochs of data and results were presented as mean ± SEM.

### RNA Extraction and qPCR

For molecular biology, mice underwent cervical dislocation, with the brain extracted and placed into a mouse brain blocker (David Kopf Instruments, Tujunga, USA). The olfactory bulbs were removed, followed by a section of tissue from Bregma 3.56 to 2.58 mm, and from Bregma −0.82 to −2.80 mm. The tissue containing prefrontal tissue was snap frozen. The posterior section was then further dissected with a scalpel to remove the amygdalar region, which was then snap frozen and stored at −80°C until use. 6 wild-type (4 male, 2 female (10.26 months ± 0.41)) and 6 App^NL-G-F^ homozygotes (3 male, 3 female (9.49 months ± 0.13)) were used for qPCR. Brain tissue of extracted regions (amygdala and prefrontal cortex) were homogenised by TissueRuptor drill (Qiagen, Manchester, UK), with ~90 mg used for RNA extraction. Instructions were followed as per the Bio-Rad Aurum Total RNA Fatty and Fibrous Tissue Kit (Bio-Rad, Hemel Hempsted, UK, Cat# 7326830) and our previously optimised protocol (Pervolaraki et al., 2018). RNA quantity and quality were confirmed by Nanodrop spectrophotometer. cDNA was generated by the iScript cDNA Synthesis Kit (Bio-Rad, UK), with 1 μg of RNA used per reaction.

Primers (Table 1) were designed using Primer3 after identifying the gene sequence on NCBI. Primers (Integrated DNA Technologies, Leuven, Belgium) were tested for specificity and run conditions optimised by PCR using whole brain cDNA. Plates were run with 10 μl per reaction, with 1 μl of cDNA, Bio-Rad SYBR Green (cat# 1725121) and 300 nM primers. Samples were run in triplicate using the protocol of 95C for 3 min, followed by 95C for 10 s, 60C for 30 s repeated 35 times. Gene expression was imaged using a Bio-Rad CFX Connect and analysed using Bio-Rad Connect Manager, and quantified using the 2^ddCt method against the housekeeping gene *Gapdh* which did not differ between the genotypes.

**Table 1.**
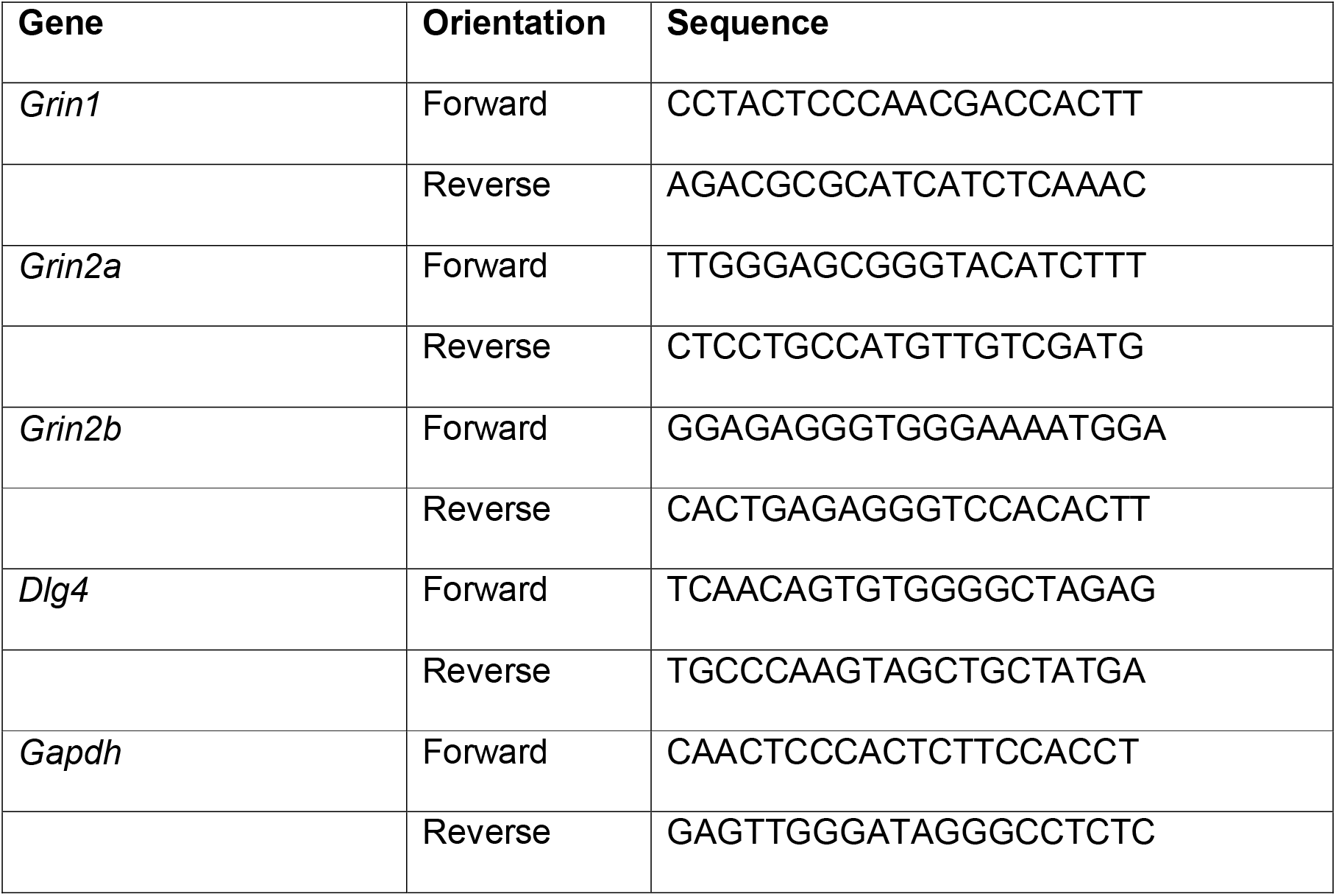

### Immunostaining

Staining for amyloid plaques can be found elsewhere (Ly et al., 2011). Tissue previously scanned by MRI was washed and immersed in 30% sucrose for >48 hrs. Tissue was cryosectioned at 30 μm, followed by immersion in 88% formic acid for 15 mins. Endogenous peroxidase activity was quenched with H_2_O_2_ for 30 mins, followed by blocking in 2% bovine serum albumin (BSA) (Thermo Fisher Scientific) for 1 hr at room temperature. Sections were then incubated with 1:500 6E10 (BioLegend, San Diego, CA, USA) overnight at 4°C, followed by 1:500 biotinylated anti-mouse (Vector Labs, Peterborough, UK) for 2 hrs at room temperature. Sections were then reacted with Vectastain ABC kit (Vector Labs) followed by diaminobenzidine (DAB) treatment (Vector Labs) prior to mounting on slides for imaging by light microscopy.

### Data Analysis

All data are expressed as mean ± standard error of the mean (SEM). To assess the variance between genotypes within a single brain structure across hemispheres, data was analysed by within subject repeated measures two-way ANOVAs or unpaired T-tests. To correct for multiple comparisons, we employed the Benjamini-Hochberg Procedure, with false discovery rate set to 0.4 (corrected P values stated). Behaviour was analysed with ANOVAs, followed by tests of simple main effects. Non-significant statistical results, particularly hemisphere comparisons, can be found in Supplemental Materials. Statistical testing and graphs were made using GraphPad Prism version 6 and SPSS v22.

## Results

### *App^NL-G-F^* KI mice show changes in anxiety-assessing tasks

To assess the potential role of Aβ in anxiety, we undertook two behavioural paradigms that are widely used to probe anxiety in rodents: the open field and the elevated plus maze. In the open field, mice were allowed to freely ambulate for 30 mins, during which we measured the distance they travelled in 5 min blocks. We observed that both *App^NL-F^* and *App^NL-G-F^* KI mice travelled a similar distance to wild-type mice (Fig. 1A; genotype F_(2, 55)_ = 2.19, p = 0.122, time block × genotype × sex F_(1, 55)_ <1, p = 0.817, genotype × sex F_(2, 55)_ = 1.21, p = 0.306), suggesting that *App* KI mice do not have any overt deficiencies in motor function. We subsequently divided the floor of the arena into an outer zone and a centre zone (see Methods) to determine whether *App* KI mice displayed a propensity to stay close to the walls (thigmotaxis) and avoid the centre. Notably, whilst wild-type and *App^NL-F^* KI mice spend a broadly similar amount of time in the outer zone, *App^NL-G-F^* KI mice spend markedly more time in the outer zone (i.e. against the walls) (Fig. 1B; genotype F_(2, 55)_ = 10.34, p <0.0001, genotype × sex F_(2, 55)_ <1, p = 0.475. Tukey’s post hoc: wild-type vs. *App^NL-G-F^* KI mice p = 0.001, *App^NL-F^* KI mice vs. *App^NL-G-F^* KI mice p = 0.001). Further suggesting an anxiogenic phenotype, *App^NL-G-F^* KI mice spent significantly less time in the centre zone (Fig. 1C; genotype F_(2, 55)_ = 7.49, p = 0.001, genotype × sex F_(2, 55)_ <1, p = 0.603. Tukey’s post hoc: wild-type vs. *App^NL-G-F^* KI mice p = 0.003, *App^NL-F^* KI mice vs. *App^NL-G-F^* KI mice p = 0.006). Finally, no differences between the genotypes were observed for entries to the outer zone or the centre zone (Supplementary Fig. 1A, 1B, see figure legend for statistics). Hence, based on both zone entries and total distance travelled (see above), there were no signs of hypo- or hyperactivity in any group.

**Figure 1.**
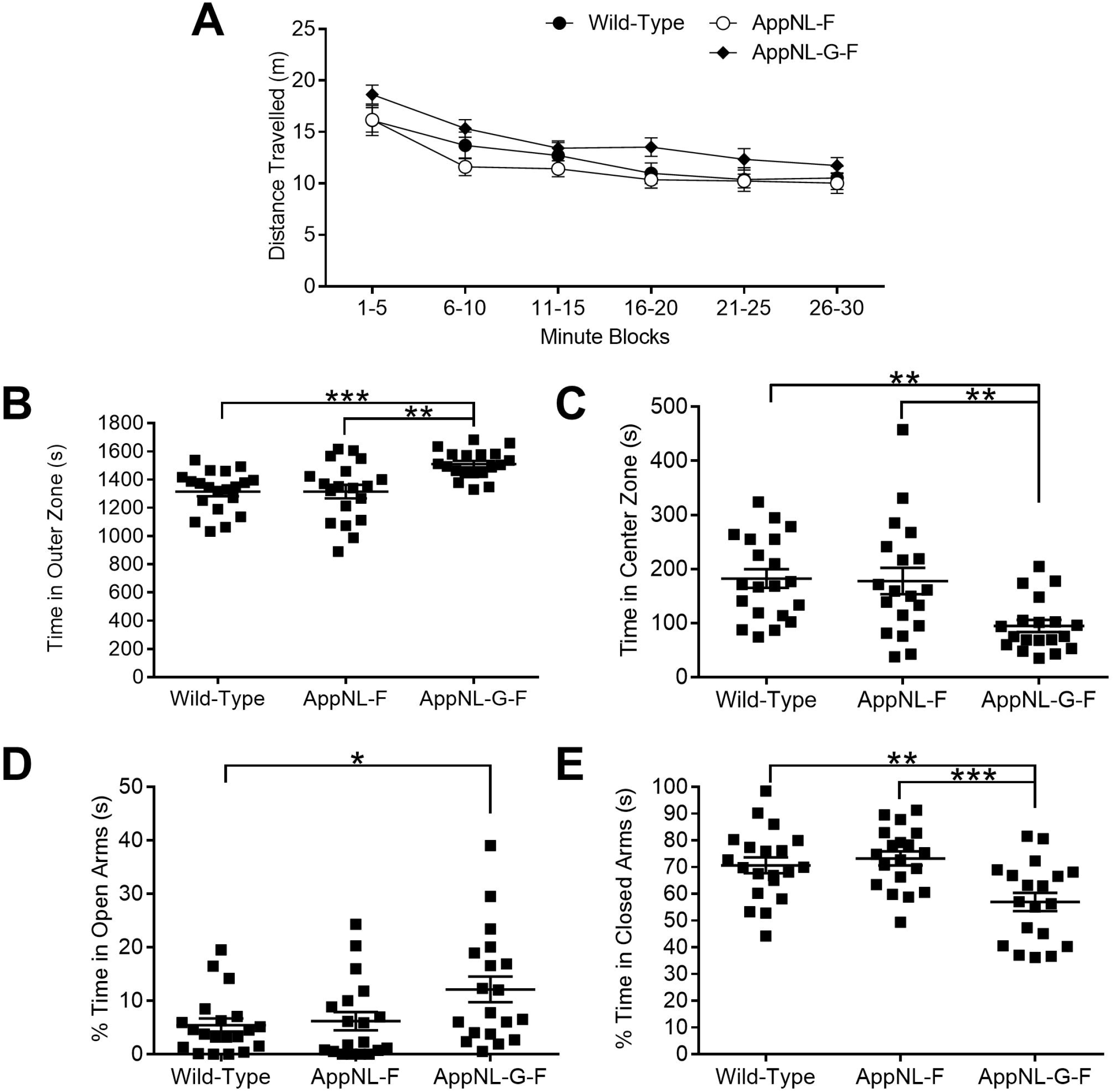
Anxiety behaviours in 8 month old *App* KI mice in the open field and elevated plus maze. **A.** Given 30 minutes free ambulation, all genotypes expressed similar amounts of locomotor activity as quantified by distance travelled. **B.** *App^NL-G-F^* KI mice displayed thigmotaxis in the open field, spending significantly more time against the walls and less time in the centre zone (**C**). Conversely, *App^NL-G-F^* KI mice spent significantly more time exploring the open arms of the elevated plus maze (**D**) and significantly less time within the closed arms (**E**), suggesting an anxiolytic profile. ***=P<0.001, **=P<0.01, *=P<0.05. Error bars = s.e.m. Wild-type n = 21, *App^NL-F^* KI n = 19, *App^NL-G-F^* KI n = 19.

We next examined whether *App^NL-G-F^* KI mice also had a similarly anxiogenic profile in the elevated plus maze (EPM). Surprisingly, *App^NL-G-F^* KI mice spent significantly more time in the EPM’s open arms (Fig. 1D; genotype F_(2, 55)_ = 4.25, p = 0.019, sex F_(1, 55)_ <1, p = 0.806, genotype × sex F_(2, 55)_ = 1.52, p = 0.229. Tukey’s post hoc: wild-type vs. *App^NL-G-F^* KI mice p = 0.032, *App^NL-F^* KI mice vs. *App^NL-G-F^* KI mice p = 0.067), and markedly less time in the EPM’s closed arms (Fig. 1E; genotype F_(2, 55)_ = 8.69, p = 0.001, sex F_(1, 55)_ <1, p = 0.664, genotype × sex F_(1, 55)_ = 1.69, p = 0.195. Tukey’s post hoc: wild-type vs. *App^NL-G-F^* KI mice p = 0.005, *App^NL-F^* KI mice vs. *App^NL-G-F^* KI mice p = 0.001). Furthermore, *App^NL-G-F^* KI mice spent markedly less time in the centre zone (see Supplementary Fig. 1C for statistics), and there were trends towards significantly more head dips made by *App^NL-G-F^* KI mice (see Supplementary Fig. 1D for statistics). As with the Open Field, the specificity of the differences is unlikely to be explained by hyper- or hypoactivity, as all genotypes travelled a similar distance within the test (see Supplementary Fig. 1E for statistics). In summary, *App^NL-G-F^* KI mice show anxiety-related that differ between experimental paradigms; anxiogenic in the Open Field, anxiolytic in the Elevated Plus Maze.

To explore whether social motivation was altered in *App* KI mice, we examined sociability in the three-chambered social approach test (Moy et al., 2004; Nadler et al., 2004) which exploits the preference of a mouse to explore a novel mouse enclosed within a wire cage compared to an identical empty cage. All genotypes showed a clear, similar preference for exploring the cage containing the novel mouse (Fig. 2A: genotype F_(2, 55)_ = 1.71, p = 0.19, genotype × discrimination between the novel mouse and empty cage F_(2, 55)_ <1, p = 0.58, genotype × sex F_(1, 55)_ = 1.98, p = 0.15).

**Figure 2.**
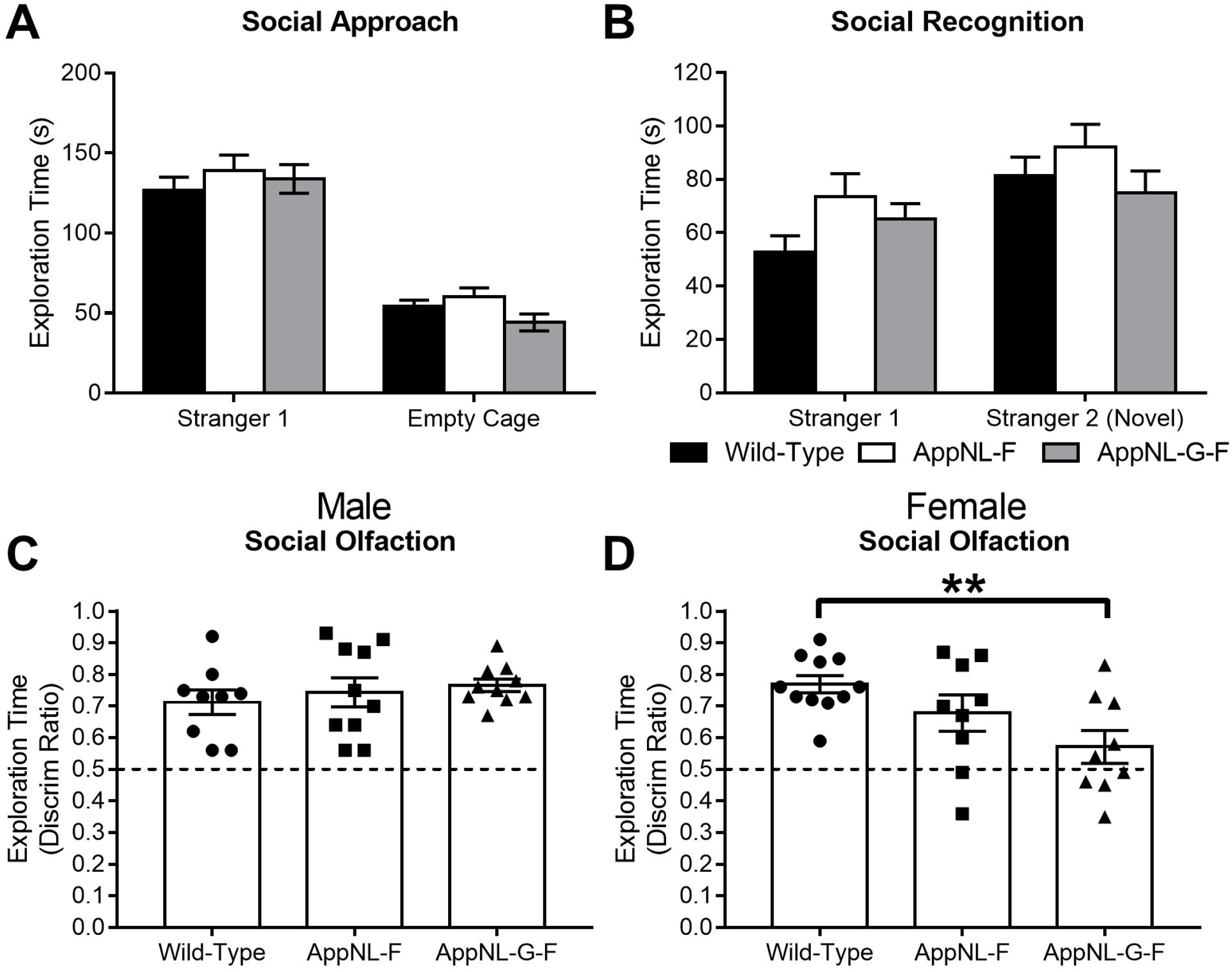
Social behaviours in *App* KI mice. **A.** All genotypes showed similar preference for exploring a novel, same-sexed novel conspecific (Stranger 1), compared to an empty cage. However, *App* KI mice displayed a weaker discrimination between the previously explored mouse (Stranger 1) and a second novel conspecific (Stranger 2) (**B**). Mice were then required to discriminate between a social smell (soiled bedding) or a non-social smell (clean bedding). Although all male genotypes show similar preference for exploring the social cue compared to male wild-types (**C**), only female *App^NL-G-F^* KI mice show significantly weaker discrimination compared to female wild-type mice (**D**). **=P<0.01. Error bars = s.e.m. Wild-type: male n = 9, female n = 11; *App^NL-F^* KI: male n = 10, female n = 9; *App^NL-G-F^* KI: male n = 10, female n = 9.

Next, we tested whether *App* KI mice were able to show social novelty recognition by preferentially exploring a second novel conspecific over a previously explored conspecific (Fig. 2B). Although we found a significant interaction between genotype and sex (F_(2, 55)_ = 3.39, p = 0.041), driven by *App^NL-G-F^* KI female mice, discrimination between the familiar and novel mouse did not differ between the genotypes (F_(1, 55)_ = 1.17, p = 0.32). Thus, only subtle deficits exist for social recognition memory in *App^NL-G-F^* but not *App^NL-F^* KI mice.

Finally, we tested whether *App* KI mice showed motivation to explore a social smell (soiled bedding) compared to a non-social smell (clean bedding). Discrimination between the social and non-social olfactory stimulus differed by sex and genotype (Supplementary Fig. 2: genotype × discrimination × sex F_(2, 55)_ = 4.31, p = 0.019, genotype × discrimination F_(2, 55)_ <1, p = 0.62, genotype × sex F_(1, 55)_ = 2.01, p = 0.14). To further investigate the source of difference, we examined social-vs-non-social smell discrimination ratios separated by sex (Fig. 2C,D). For males, all genotypes showed discrimination for the social stimulus (one-way ANOVA, F_(2, 26)_ <1). However, female *App^NL-G-F^* KI mice displayed significantly weaker discrimination between the social and non-social olfactory stimulus compared to female wild-types (one-way ANOVA, F_(2, 26)_ = 4.91, p = 0.016. Dunnett’s post-hoc test wild-type vs. *App^NL-G-F^* KI p = 0.008).

### *App^NL-G-F^* KI mice have microstructural changes in the prefrontal cortex and hippocampus

Given that behavioural changes in the open field, elevated plus maze and social olfaction test are exclusively restricted to *App^NL-G-F^* KI mice, we decided to take this genotype forward for further analysis to determine the biological mechanisms that may explain these impairments. Our approach was to examine the integrity of brain regions associated with both social and anxiety behaviours (see Methods). These centred upon the prefrontal cortex (OFC and ACC), the hippocampus (anterior and posterior) and the amygdala (including the basolateral (BLA)). To derive quantitative measures of DTI, we examined FA and MD (examining diffusion across the λ_1_, λ_2_ and λ_3_ vectors), followed by AD and RD (to determine preferential diffusion along the λ_1_, or λ_2_ and λ_3_ vectors, respectively).

Given the amygdala has been widely associated with both social and anxiety behaviours (Adolphs, 2010; Davis, 1992; Davis et al., 1994), we segmented the whole amygdalar region, in anterior and posterior planes, and separately, the basolateral nuclei (BLA), to determine whether structural alterations may explain the behavioural changes. For all measures (FA, MD, AD and RD), in both the anterior and posterior amygdala, plus the BLA, we did not observe any significant changes in tissue diffusion properties (Supplementary Table 1 for non-significant statistics and Supplementary Fig. 3).

Ventral hippocampal regions are associated with unconditioned anxiety as tested by OF and EPM tasks (Bannerman et al., 2004), whilst dorsal and ventral regions are increasingly being associated with social recognition (Hitti and Siegelbaum, 2014; Okuyama et al., 2016). We therefore next examined the microstructure of the hippocampus along the anterior to posterior axis. FA of the anterior (Fig. 3A) and posterior (Fig. 3B) hippocampus did not differ between wild-types (Supplementary Table 2 for non-significant statistics). However, whilst MD in the anterior hippocampus (Fig. 3C) was similar between the genotypes F_(1, 14)_ <1, p = 0.263), MD was significantly higher in the posterior hippocampus (Fig. 3D) of *App^NL-G-F^* KI mice (F_(1, 14)_ = 12.18, p = 0.011). For AD, *App^NL-G-F^* KI mice had significantly increased diffusion in the anterior hippocampus (Fig. 3E F_(1, 14)_ = 7.55, p = 0.047) but not in the posterior hippocampus (Fig. 3F F_(1, 14)_ = 4.45, p = 0.068). RD was significantly higher in *App^NL-G-F^* KI mice in the posterior hippocampus (Fig. 3G F_(1, 14)_ = 18.13, p = 0.005) but not the anterior hippocampus (Fig. 3H F_(1, 14)_ = 1.52, p = 0.142). In summary, diffusion changes in the hippocampus were not marked, with significant changes only in anterior hippocampal AD, and posterior hippocampal RD.

**Figure 3.**
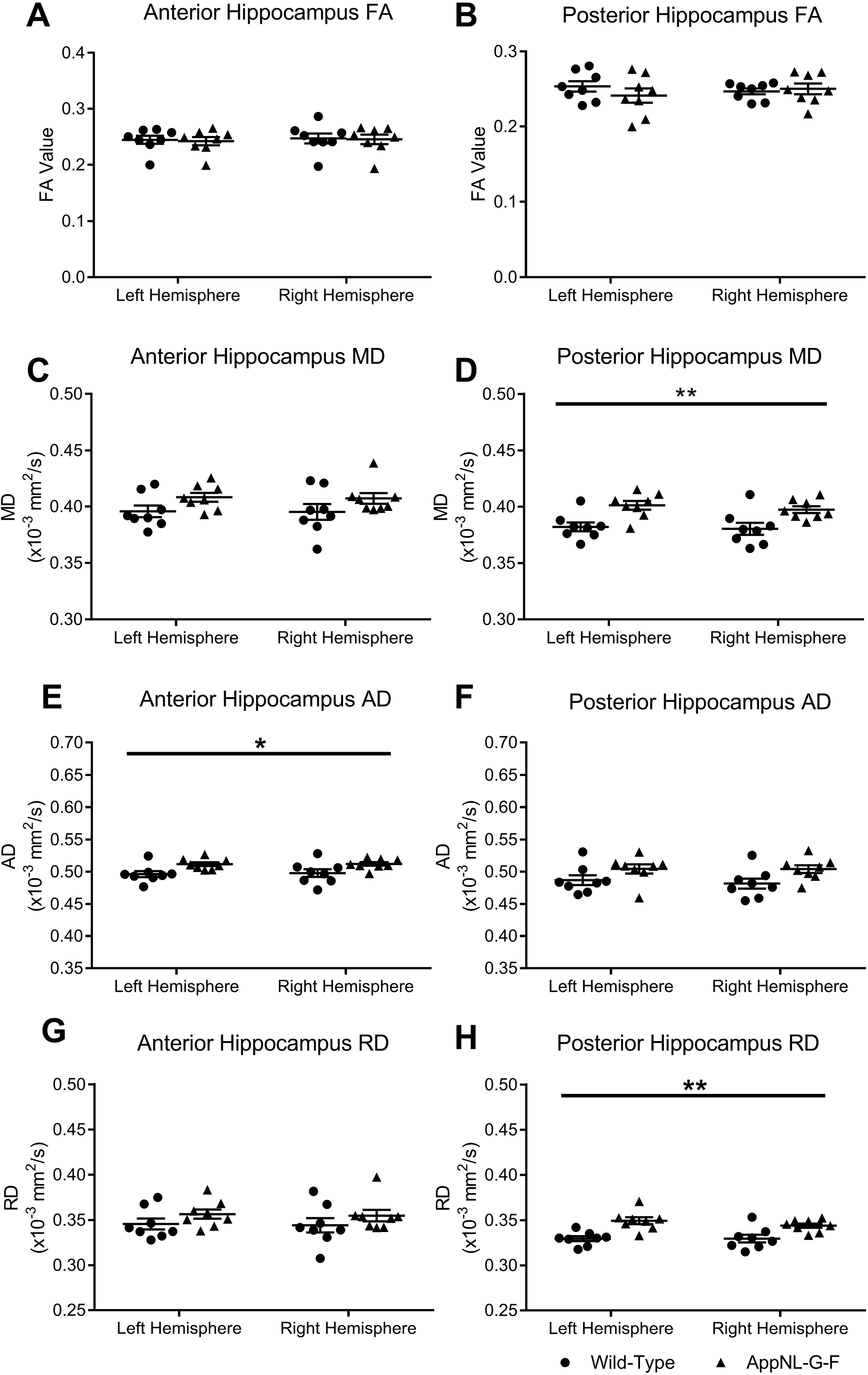
Alterations in fractional anisotropy (FA) and mean diffusivity (MD) in the hippocampus. DTI images of the hippocampus was segmented at two regions; anterior (Bregma −1.94 mm) and posterior (Bregma −3.28 mm). FA was not significantly altered between the wild-type and *App^NL-G-F^* KI mice in the anterior (**A**) and posterior (**B**) hippocampus. MD of the anterior hippocampus did not differ between the genotypes (**C**), however MD in the posterior hippocampus was significantly higher in *App^NL-G-F^* KI mice (**D**). Diffusivity was further characterised by examining axial (AD) and radial (RD) diffusivity. AD was significantly increased in the anterior hippocampus of *App^NL-G-F^* KI mice (**E**), but not in the posterior hippocampus (**F**). RD did not differ between the genotypes in the anterior hippocampus, but *App^NL-G-F^* KI mice had significantly increased RD posterior hippocampus compared to wild-types. **=P<0.01, *=P<0.05. Error bars = s.e.m. Wild-type n = 8, *App^NL-G-F^* KI n = 8.

We next segmented two prefrontal cortical regions: the OFC and the ACC. In addition to roles for OFC and ACC regions in anxiety and social behaviours, resting state functional MRI (rsfMRI) has shown that connectivity of the medial prefrontal cortex to other regions is abnormal in *App^NL-G-F^* KI mice, with ACC being the most altered region (Latif-Hernandez et al., 2017). However, it is currently unknown as to whether structural changes in the ACC contributed to the rsfMRI result. In the OFC of *App^NL-G-F^* KI mice, there were no significant differences in FA (Fig. 4A; genotype (F_(1, 14)_ = 4.51, p = 0.063), however MD was significantly increased (Fig. 4B; genotype (F_(1, 14)_ = 8.61, p = 0.026). Similarly, for the ACC, FA was similar between the genotypes (Fig. 4C; genotype (t_(14)_ <1, p = 0.37) but MD was significantly increased (Fig. 4D; genotype (t_(14)_ = 3.13, p = 0.021). We further explored prefrontal cortical changes by quantifying AD and RD. In the OFC, both AD and RD were significantly increased in *App^NL-G-F^* KI mice (F_(1, 14)_ = 8.34, p = 0.032 (Fig. 4E) and F_(1, 14)_ = 8.23, p = 0.042 (Fig. 4F), respectively). AD and RD were also significantly increased in the ACC of *App^NL-G-F^* KI mice (t_(14)_ = 2.88, p = 0.037 (Fig. 4G) and t_(14)_ = 3.19, p = 0.016 (Fig. 4H). In summary, for both OFC and ACC, FA was not significantly altered, but MD, AD, and RD were all increased in *App^NL-G-F^* KI mice. Significant increases in three of these DTI measures likely reflects widespread pathology in the OFC and ACC of *App^NL-G-F^* KI mice. The DTI-related alterations in these prefontal regions notably contrasted with an absence of any alterations in DTI measures in the amygdala.

**Figure 4.**
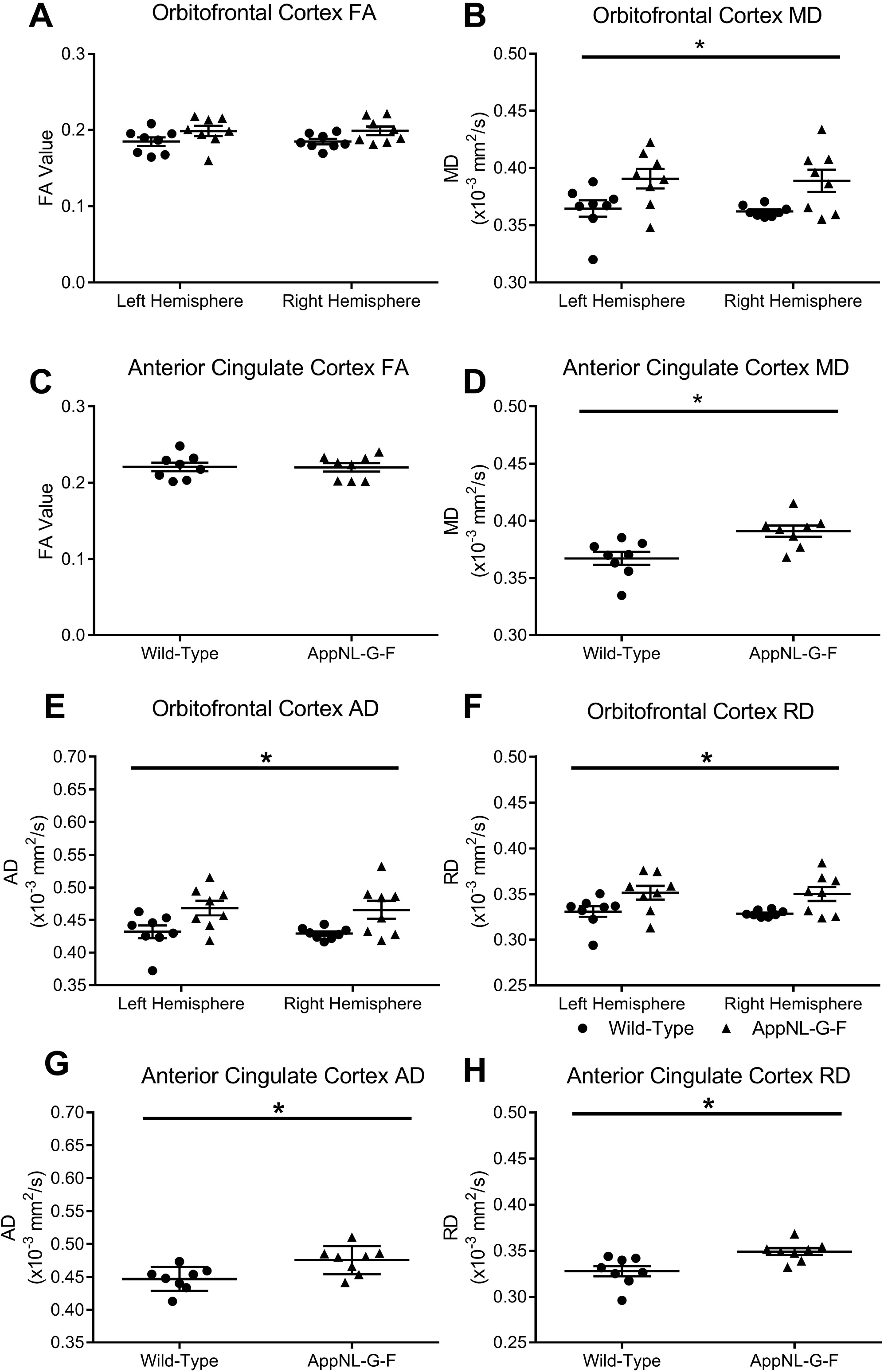
Alterations in fractional anisotropy (FA) and mean diffusivity (MD) in the orbitofrontal cortex and anterior cingulate cortex of the prefrontal cortex. DTI images were segmented for the orbitofrontal cortex at Bregma +2.58 mm and for the anterior cingulate cortex at Bregma +1.18 mm. FA was not significantly altered between the wild-type and *App^NL-G-F^* KI mice in the orbitofrontal cortex (**A**), however MD was significantly higher for *App^NL-G-F^* KI mice (**B**). Similarly, FA was did not differ between the genotypes for the anterior cingulate cortex (**C**), but MD was significantly higher in *App^NL-G-F^* KI mice (**D**). Axial (AD) and radial (RD) diffusivity was then analysed. In *App^NL-G-F^* KI mice, AD (**E**) and RD (**F**) were both significantly increased in the orbitofrontal cortex. AD (**G**) and RD (**H**) were also both increased in *App^NL-G-F^* KI mice in the anterior cingulate cortex. *=P<0.05. Error bars = s.e.m. Wild-type n = 8, *App^NL-G-F^* KI n = 8.

Finally, we performed amyloid plaque staining on the corresponding tissue sections that were analysed for DTI. By 9 months of age in *App^NL-G-F^* KI mice, the OFC, ACC, amygdala and hippocampus all exhibit substantial plaque load (Supplementary Fig. 4), indicating that the lack of DTI changes in the amygdala are not due to an absence of amyloid plaques.

### Alterations in prefrontal cortical NMDA-dependent gamma oscillations in *App^NL-G-F^* KI mice

Although some microstructure changes were detected in the hippocampus of *App^NL-G-F^* KI mice (anterior hippocampal AD and posterior hippocampal RD), these were relatively inconsistent compared to the alterations observed within the prefrontal cortex. Another study found the ACC as being the most significantly altered brain region in *App^NL-G-F^* KI mice as detected by rsfMRI (Latif-Hernandez et al., 2017). This suggests that substantial changes are occurring in the ACC in *App^NL-G-F^* KI mice, as broadly consistent with ACC amyloid-beta load being high in MCI, and predicting faster conversion to AD (Okello et al., 2009). As such, and given the importance of the ACC and amygdala to anxiety and social behaviours, we decided to contrast these two regions to examine whether altered network physiology might accompany DTI-derived microstructural alterations.

Gamma oscillations have been widely associated with roles in learning and memory (Tallon-Baudry et al., 1998) and attention (Fries et al., 2001), whilst coherence between brain regions has been demonstrated to facilitate information transfer (Buzsaki and Wang, 2012). Specifically in the prefrontal cortex, gamma oscillations have been shown to be important for social behaviour (Cao et al., 2018) and anxiety behaviours (Dzirasa et al., 2011; Stujenske et al., 2014). Gamma disruption has been demonstrated in other AD mouse models, such as the APP/PS1 (Klein et al., 2016), the TAS10 overexpression model (Driver et al., 2007) and in the entorhinal cortex of *App^NL-G-F^* KI mice (Nakazono et al., 2017), and these impairments occur relatively early in Aβ-driven pathological processes. Thus, gamma oscillations represent a useful target for exploring network-level sequelae of AD-related pathology, relevant to social and anxiety-related behaviour.

We generated gamma oscillations in brain slices using kainate and tested their dependency on NMDA receptors, which have previously been shown to modulate peak frequency through inhibitory postsynaptic currents (Carlen et al., 2012; McNally et al., 2011) by modifying recruitment of different interneuron subpopulations (Middleton et al., 2008). Within the BLA of the amygdala, we found that in wild-types, peak amplitude and frequency of gamma oscillations were unaffected by the application of the broad spectrum antagonist NMDA receptor antagonist CPP (Fig. 5Aiii; t_(14)_ <1, p = 0.991 and t_(14)_ <1, p = 0.656, respectively). We observed similar results in the *App^NL-G-F^* KI mice, with peak amplitude and frequency unaffected by NMDA antagonism (Fig. 5Aiv; t_(14)_ <1, p = 0.951 and t_(14)_ <1, p = 0.709, respectively). Together, this suggests that gamma oscillations in the BLA in *App^NL-G-F^* KI mice are unaffected by Aβ deposition.

**Figure 5.**
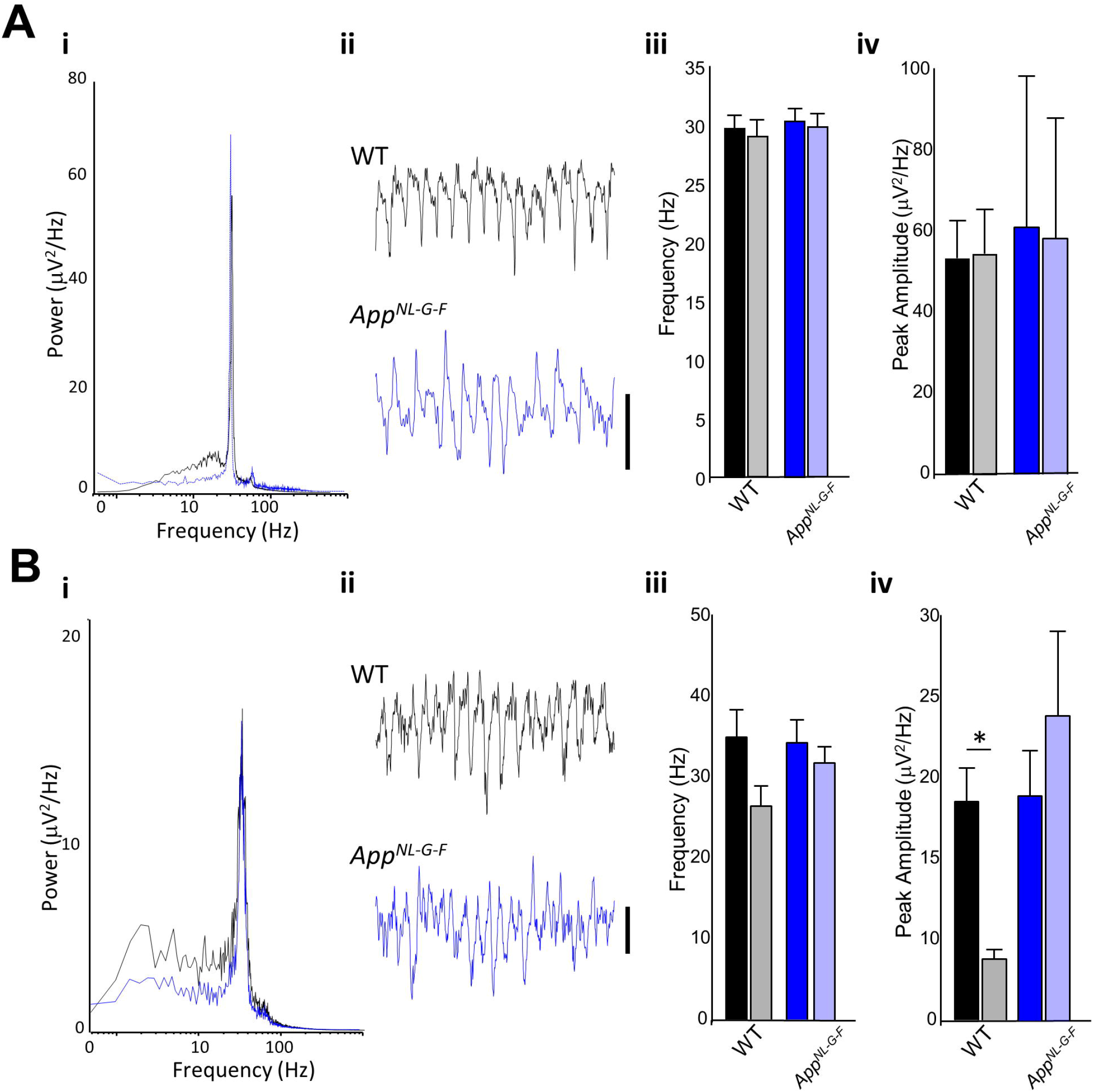
Gamma oscillations in the amygdala and anterior cingulate cortex. **A. i.** Pooled power spectrum of gamma oscillatory activity in the basolateral amygdala in WT (black) and *App^NL-G-F^* (blue) KI mice. **ii.** Example traces showing 500 ms of gamma oscillatory activity in WT (black) and *App^NL-G-F^* (blue) KI mice. Scale bar 50 mV. **iii.** Graph showing the effect of NMDA receptor antagonism (20 mM CPP) on the frequency of basolateral amygdala gamma oscillations. **iv.** Graph showing the effect of NMDA receptor antagonism (20 mM CPP) on the amplitude of basolateral amygdala gamma oscillations. **B. i.** Pooled power spectrum of gamma oscillatory activity in the anterior cingulate cortex in WT (black) and *App^NL-G-F^* (blue) KI mice. **ii.** Example traces showing 500 ms of gamma oscillatory activity in WT (black) and *App^NL-G-F^* (blue) KI mice. Scale bar 20 mV. **iii.** Graph showing the effect of NMDA receptor antagonism (20 mM CPP) on the frequency of anterior cingulate cortex gamma oscillations. **iv.** Graph showing the effect of NMDA receptor antagonism (20 mM CPP) on the amplitude of anterior cingulate cortex gamma oscillations. *=P<0.05. Error bars = s.e.m. Wild-type n = 5, *App^NL-G-F^* KI n = 5.

Next, we studied gamma oscillations within the ACC of the prefrontal cortex. In wild-types, the frequency of gamma oscillations was unaffected by CPP (Fig. 5Biii; t_(14)_ = 1.91, p = 0.086). However, gamma peak amplitude was significantly reduced by CPP application (Fig. 5Biv; t_(14)_ = 2.96, p = 0.014), suggesting that, as expected (e.g. (Carlen et al., 2012)) ACC gamma oscillations normally require NMDA receptors in the wild-type conditions. In contrast, in *App^NL-G-F^* KI mice, gamma amplitude (and frequency) were unaltered by NMDA receptor antagonism (t_(14)_ <1, p = 0.404; t_(14)_ <1, p = 0.430 respectively), suggesting that ACC gamma oscillations in *App^NL-G-F^* KI mice have lost their dependency upon NMDA receptors. In summary, the contribution of NMDA receptors to gamma oscillatory mechanisms appeared abnormally reduced in the prefrontal cortex.

To further understand this pattern of NMDA receptor-related alterations in gamma in the prefrontal cortex but not BLA amygdala, we next analysed the mRNA expression of synaptic genes including NMDA receptors and pre and postsynaptic receptors relating to NMDA receptor function. Within the amygdala, although we generally observed higher gene expression in *App^NL-G-F^* KI mice (Fig. 6A), no significant genotypic differences were observed for *Dlg4* (t_(10)_ = 2.14, p = 0.058), *Grin1* (t_(10)_ = 1.97, p = 0.078), *Grin2a* (t_(10)_ = 2.16, p = 0.056), *Grin2b* (t_(10)_ <1, p = 0.455) or *Stxbp1* (t_(10)_ = 1.27, p = 0.234). We next examined the mRNA expression in the frontal cortex (Fig. 6B). We did not observe any significant differences between wild-types and *App^NL-G-F^* KI mice for the genes *Dlg4* (t_(10)_ <1, p = 0.420), *Grin1* (t_(10)_ <1, p = 0.780), and *Grin2a* (t_(10)_ <1, p = 0.731). However, the expression of *Grin2b* [protein: 2B subunit of NMDA-receptor] (t_(10)_ = 2.59, p = 0.027) and *Stxbp1* [protein: Munc18-1] (t_(10)_ = 2.66, p = 0.024) was significantly reduced in *App^NL-G-F^* KI mice. Together, the change in the dependency of NMDA receptor mediated gamma rhythms in the ACC of *App^NL-G-F^* KI mice could be related to reduced *Grin2b* or presynaptic release through Munc18-1.

**Figure 6.**
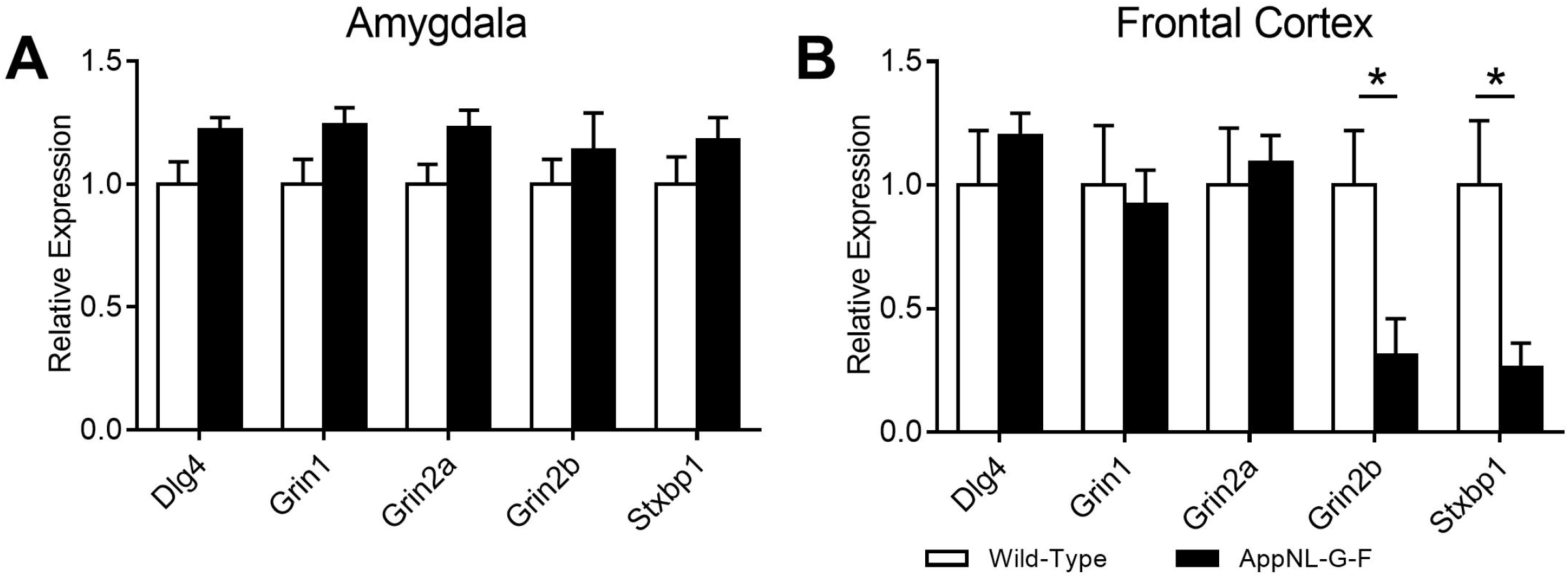
Gene expression within the frontal cortex and amygdala. **A.** Although *App^NL-G-F^* KI mice generally had higher expression of our selected genes within the amygdala, these were not significantly different to wild-types. **B.** Within the frontal cortex, expression of both *Grin2b* and *Stxbp1* (Munc18-1) were significantly reduced in *App^NL-G-F^* KI mice. *=P<0.05. Error bars = s.e.m. Wild-type n = 6, *App^NL-G-F^* KI n = 6.

## Discussion

Within the current study, we have further characterised the behavioural profile of the new generation of *App* KI mice. When tested at 8 months, *App^NL-G-F^*, but not *App^NL-F^*, knock-in mice show changes in behaviour relative to controls. *App^NL-G-F^* KI mice exhibited subtle deficits in social recognition and social olfactory discrimination, but not in sociability. In anxiety-assessing tasks, *App^NL-G-F^* KI mice exhibited task-dependent changes: 1) In the open field, they showed increased thigmotaxis, and reduced centre time, i.e. an ostensibly-anxiogenic profile; 2) in the Elevated Plus Maze (EPM), they showed reduced closed-arm, and increased open-arm, time, i.e. an ostensibly-anxiolytic profile. We discuss this seemingly contradictory profile further below. Briefly, one view is that these anxiety-related changes in *App^NL-G-F^* knock-in mice, being contradictory, likely do not reflect a simple shift towards one or other extreme of emotionality. Rather, their combined ostensibly-anxiogenic open field, yet ostensibly-anxiolytic EPM, profile could hint at altered mechanisms affecting decision-making (e.g. ‘disinhibition’ in the EPM). If so, this idea would predict mild/absent amygdalar changes combined with marked prefrontal cortical changes. Consistent with this idea, alterations in microstructure, glutamatergic-dependent gamma oscillations, and glutamatergic gene expression were all observed in the prefrontal cortex, but not the amygdala, of *App^NL-G-F^* knock-in mice. Specifically, in *App^NL-G-F^* mice: DTI detected increases in mean, radial and axial diffusivity in the Orbitofrontal and Anterior Cingulate cortices; slice recordings showed that ACC prefrontal gamma mechanisms were unusual in being NMDA-R independent; finally, likely relatedly, qPCR showed reduced expression of *Grin2b* [protein: 2B subunit of NMDA-receptor] in prefrontal cortex. We speculate that abnormally NMDA-R independent ACC gamma could reflect compensatory changes following reduced NMDA-R expression.

*App^NL-F^* KI and *App^NL-G-F^* KI mice enable researchers to dissociate the amyloidogenic effects of Aβ deposition. At ~9 months of age, *App^NL-G-F^* KI mice have ~100 times more insoluble Aβ_42_ than *App^NL-F^* KI mice, but the same amount of soluble Aβ_42_ (Saito et al., 2014). Accordingly, any dysfunction found in *App^NL-F^* KI *but not App^NL-G-F^* KI mice of ~9 months age, as here, can reasonably be attributed as the sequelae of specifically insoluble Aβ_42_ aggregation. We therefore argue that specifically insoluble Aβ_42_ aggregation drove the behavioural changes that were seen in *App^NL-G-F^* KI but not *App^NL-F^* KI mice: higher thigmotaxis and reduced centre-time in the open field, more time in the open arms, and less in the centre of the elevated plus maze, and deficits in the social olfaction test.

Within the current study, we observed a conflicting phenotype between increased thigmotaxis in the open field (anxiogenic) and increased time in the open arms in the elevated plus maze (anxiolytic). Others have examined the anxiety profile in *App^NL-G-F^* KI mice using the open field and found that 6 month old mice spent more time in the open field centre zone (Latif-Hernandez et al., 2017). However, others have found no difference in time spent in the centre of an open field for 6 month old *App^NL-G-F^* KI mice (Whyte et al., 2018). Latif-Hernandez et al. (2017) and Whyte et al. (2018) did not specifically quantify thigmotaxis in the open field, limiting comparisons to our study. Interestingly, locomotion within the open field appears to vary by age. Distance travelled in the open field is significantly greater in 6 month old *App^NL-G-F^* KI mice, whilst at 8 months (the current study) and 10 months of age, there were no significant differences (Latif-Hernandez et al., 2017; Whyte et al., 2018), suggesting hyperactivity is only detectable in younger mice.

Like our study, Latif-Hernandez et al. (2017) also used the elevated plus maze to explore anxiety behaviours in *App^NL-G-F^* KI mice. However, unlike us, they compared *App^NL-G-F^* KI mice behaviour to that of *App^NL^* KI mice only, and not to control WT mice. They found *App^NL-G-F^* KI mice spent more time in the open arms at 6 months of age (Latif-Hernandez et al., 2017), which we replicated at 8 months of age. A very recent study has also found that 6-9 month old *App^NL-G-F^* KI mice spent more time in the elevated plus maze open arms (Sakakibara et al., 2018). Together with our study, then, three independent studies suggest that *App^NL-G-F^* KI mice display an anxiolytic profile in the elevated plus maze. The ostensible contradiction between our results for the *App^NL-G-F^* KI mice in the open field (higher thigmotaxis) and elevated plus maze (lower centre time, higher open arm time) is curious but not unique. Tg2576 mice also show open field thigmotaxis yet increased time in the elevated plus maze open arms (Lalonde et al., 2003). It is possible that the increased time spent in the open arms reflects a disinhibition phenotype. Disinhibition is a well-established, albeit less common, AD phenotype (Chung and Cummings, 2000; Hart et al., 2003), which could be manifested within the current study as a failure to inhibit the choice to enter the open arm. Given the lack of specific data to speak to a disinhibition hypothesis, it is clear that future anxiety testing in *App^NL-G-F^* KI mice will require multiple paradigms to clearly delineate anxiety from disinhibition. A benefit of our study is that we are the first to behaviourally compare *App^NL-F^* KI and *App^NL-G-F^* KI mice beyond the original Saito (2014) paper. Regarding the open field and elevated plus maze, genotypic differences were only found in *App^NL-G-F^* KI mice, suggesting behavioural modifications were primarily driven by insoluble Aβ. Currently, aged *App^NL-F^* KI mice have not been tested for anxiety behaviours, but it would be interesting to see if as these mice age, and plaque pathology increases, their behavioural phenotype becomes more like the *App^NL-G-F^* KI genotype.

Several other AD mouse models exhibit impairments in social behaviour, including APP_swe_/PS1 (Filali et al., 2011) and Tg2576 mice (Deacon et al., 2009). Latif-Hernandez et al. (2017) examined social behaviours in *App^NL-G-F^* KI mice and found that, although not significant, the discrimination ratio for exploring a novel mouse compared to an empty cage, and for comparing between the previously explore mouse and a second novel mouse, had fallen to chance by 10 months of age (Latif-Hernandez et al., 2017). We found that *App^NL-F^* KI and *App^NL-G-F^* KI mice show robust preference for exploring a novel conspecific compared to an empty cage. We did, however, find that social preference as measured by discriminating a social odour cue was impaired in female *App^NL-G-F^* KI mice. A key difference between our study and Latif-Hernandez et al. (2017) is that they used only female mice. Our study, with mixed-sex groups, suggests that females show a greater impairment in social behaviour than males, which will be an important consideration for future studies. Such female-linked deficits are broadly consistent with human AD literature: in humans, even adjusting for age and education, women are more likely to be diagnosed with AD (but not vascular dementia) than men (Andersen et al., 1999; Baum, 2005).

The ACC and amygdala are brain regions well established with mediating anxiety/fear and social behaviours (Davidson, 2002; Davis, 1992; Davis et al., 1994; Etkin et al., 2011). Additionally, the prefrontal cortex has a well-established link to behavioural decision-making, especially for affective stimuli (Dias et al., 1996). This may relate to our proposed ‘disinhibition’ explanation for the dichotomy in our two anxiety tasks, highlighting the prefrontal cortex as the critical brain region in *App^NL-G-F^* KI mice. Furthermore, these areas are also susceptible to degeneration in early AD pathology (Huang et al., 2002; Poulin et al., 2011; Scheff and Price, 2001). Other transgenic AD mouse models, notably the APP/PS1 mouse, exhibit early amygdala pathology (Guo et al., 2017; Lin et al., 2015), whilst Aβ deposition in the PDAPP mouse begins in the cingulate cortex (Schenk et al., 1999). With these findings in mind, we hypothesised that structural and functional alterations within the amygdala and medial prefrontal cortex could partly explain behavioural impairments. As it turned out, we found little evidence for neurodegeneration-linked changes in the amygdala, but did find such changes in the prefrontal cortex.

Our analysis of microstructural integrity using DTI revealed that the prefrontal cortex of *App^NL-G-F^* KI mice, including the OFC and ACC, was substantially altered, and the functional consequences of these changes to the ACC was that gamma oscillations lost their dependency upon NMDA receptors, notably GluN2B receptors. Thus far, there have been limited investigations into the biological pathways altered in *App^NL-G-F^* KI mice. Using rsfMRI, Latif-Hernandez et al. (2017) found that the cingulate cortex was the most significantly altered brain region, with no visible impairment in the amygdala. Although further work is required to definitively describe the physiological pathways that explain the altered behaviour in *App^NL-G-F^* KI mice, our findings present an important step in this process. Although other brain regions likely contribute to changes in anxiety and social behaviours (gamma oscillations have also been found altered in the medial entorhinal cortex of *App^NL-G-F^* KI mice (Nakazono et al., 2017)), the prefrontal cortex could represent a key region. Neural oscillations in the medial prefrontal cortex have clear links to anxiety behaviours, both in the open field and elevated plus maze (Adhikari et al., 2010), and medial prefrontal cortex gamma is required for the expression of social novelty (Cao et al., 2018). Additionally, the loss of GluN2B or its phosphorylation impairs social behaviour (Jacobs et al., 2015; Wang et al., 2011) and increases anxiety in the open field (Hanson et al., 2014) and the elevated plus maze (Delawary et al., 2010). Furthermore, NMDA receptors are necessary for gamma mediated by the goblet cell interneurons within the entorhinal cortex (Middleton et al., 2008), with specific antagonism of GluN2B significantly reducing hippocampal gamma power (Hanson et al., 2013). Together, insoluble Aβ overexpression drives prefrontal cortical alterations through gamma oscillatory impairments by reduced GluN2B expression. The lack of significant changes to gamma, gene expression and microstructure in the amygdala suggests that it may not undergo pathological damage at the same rate as other regions. This indicates that social and anxiety behavioural impairments in *App^NL-G-F^* KI mice are driven by regions that do not include the amygdala, although further work will be required to corroborate this.

The findings presented herein show a clear dichotomy of anxiety behaviours between two different paradigms (anxiolytic-like in the elevated plus maze vs anxiogenic-like in the open field). We have argued that this pattern may be linked to degeneration in neural regions related to decision-making, notably the medial prefrontal cortex, rather than to emotionality *per se.* Contrasted to *App^NL-F^* KI mice, only *App^NL-G-F^* KI mice exhibited any change in anxiety behaviours, suggesting that plaque-based Aβ is responsible for these effects. We further postulate that microstructural integrity, gamma oscillatory function and *Grin2b* expression within the prefrontal cortex may, in part, explain these behavioural changes. Our data continues to highlight the importance of using AD models that do not have the confound of APP overexpression. Further studies will be required to continue to refine the mechanisms that explain anxiety impairments in *App^NL-G-F^* KI mice.

## Acknowledgements

This work was supported an Alzheimer’s Society (UK) Fellowship (AS-JF-15-008) to JD, a BBSRC grant (BB/M008975/1) to CL, and a Wellcome Trust (UK) grant to MAW.

## References

Adhikari, A., et al., 2010. Synchronized activity between the ventral hippocampus and the medial prefrontal cortex during anxiety. Neuron. 65, 257–69.

Adolphs, R., 2010. What does the amygdala contribute to social cognition? Ann N Y Acad Sci. 1191, 42–61.

Andersen, K., et al., 1999. Gender differences in the incidence of AD and vascular dementia: The EURODEM Studies. EURODEM Incidence Research Group. Neurology. 53, 1992–7.

Bannerman, D. M., et al., 2004. Regional dissociations within the hippocampus--memory and anxiety. Neurosci Biobehav Rev. 28, 273–83.

Baum, L. W., 2005. Sex, hormones, and Alzheimer’s disease. J Gerontol A Biol Sci Med Sci. 60, 736–43.

Bennett, D. A., et al., 2006. The effect of social networks on the relation between Alzheimer’s disease pathology and level of cognitive function in old people: a longitudinal cohort study. Lancet Neurol. 5, 406–12.

Bloom, G. S., 2014. Amyloid-beta and tau: the trigger and bullet in Alzheimer disease pathogenesis. JAMA Neurol. 71, 505–8.

Buzsaki, G., Wang, X. J., 2012. Mechanisms of gamma oscillations. Annu Rev Neurosci. 35, 203–25.

Cao, W., et al., 2018. Gamma Oscillation Dysfunction in mPFC Leads to Social Deficits in Neuroligin 3 R451C Knockin Mice. Neuron. 97, 1253–1260 e7.

Carlen, M., et al., 2012. A critical role for NMDA receptors in parvalbumin interneurons for gamma rhythm induction and behavior. Mol Psychiatry. 17, 537–48.

Chung, J. A., Cummings, J. L., 2000. Neurobehavioral and neuropsychiatric symptoms in Alzheimer’s disease: characteristics and treatment. Neurol Clin. 18, 829–46.

Dachtler, J., et al., 2014. Deletion of alpha-neurexin II results in autism-related behaviors in mice. Transl Psychiatry. 4, e484.

Davidson, R. J., 2002. Anxiety and affective style: role of prefrontal cortex and amygdala. Biol Psychiatry. 51, 68–80.

Davis, M., 1992. The role of the amygdala in fear and anxiety. Annu Rev Neurosci. 15, 353–75.

Davis, M., et al., 1994. Neurotransmission in the rat amygdala related to fear and anxiety. Trends Neurosci. 17, 208–14.

Deacon, R. M., et al., 2009. Aged Tg2576 mice are impaired on social memory and open field habituation tests. Behav Brain Res. 197, 466–8.

Delawary, M., et al., 2010. NMDAR2B tyrosine phosphorylation regulates anxiety-like behavior and CRF expression in the amygdala. Mol Brain. 3, 37.

Devanand, D. P., et al., 2007. Hippocampal and entorhinal atrophy in mild cognitive impairment: prediction of Alzheimer disease. Neurology. 68, 828–36.

Dias, R., et al., 1996. Dissociation in prefrontal cortex of affective and attentional shifts. Nature. 380, 69–72.

Driver, J. E., et al., 2007. Impairment of hippocampal gamma-frequency oscillations in vitro in mice overexpressing human amyloid precursor protein (APP). Eur J Neurosci. 26, 1280–8.

Dzirasa, K., et al., 2011. Impaired limbic gamma oscillatory synchrony during anxiety-related behavior in a genetic mouse model of bipolar mania. J Neurosci. 31, 6449–56.

Etkin, A., et al., 2011. Emotional processing in anterior cingulate and medial prefrontal cortex. Trends Cogn Sci. 15, 85–93.

Ferretti, L., et al., 2001. Anxiety and Alzheimer’s disease. J Geriatr Psychiatry Neurol. 14, 52–8.

Filali, M., et al., 2011. Anomalies in social behaviors and exploratory activities in an APPswe/PS1 mouse model of Alzheimer’s disease. Physiol Behav. 104, 880–5.

Franklin, K., Paxinos, G., 2008. The Mouse Brain in Stereotaxic Coordinates. Academic Press.

Fries, P., et al., 2001. Modulation of oscillatory neuronal synchronization by selective visual attention. Science. 291, 1560–3.

Gallagher, D., et al., 2011. Anxiety and behavioural disturbance as markers of prodromal Alzheimer’s disease in patients with mild cognitive impairment. Int J Geriatr Psychiatry. 26, 166–72.

Guo, C., et al., 2017. Early-stage reduction of the dendritic complexity in basolateral amygdala of a transgenic mouse model of Alzheimer’s disease. Biochem Biophys Res Commun. 486, 679–685.

Hanson, J. E., et al., 2014. Chronic GluN2B antagonism disrupts behavior in wild-type mice without protecting against synapse loss or memory impairment in Alzheimer’s disease mouse models. J Neurosci. 34, 8277–88.

Hanson, J. E., et al., 2013. GluN2B antagonism affects interneurons and leads to immediate and persistent changes in synaptic plasticity, oscillations, and behavior. Neuropsychopharmacology. 38, 1221–33.

Hart, D. J., et al., 2003. A retrospective study of the behavioural and psychological symptoms of mid and late phase Alzheimer’s disease. Int J Geriatr Psychiatry. 18, 1037–42.

Hitti, F. L., Siegelbaum, S. A., 2014. The hippocampal CA2 region is essential for social memory. Nature. 508, 88–92.

Huang, C., et al., 2002. Cingulate cortex hypoperfusion predicts Alzheimer’s disease in mild cognitive impairment. BMC Neurol. 2, 9.

Jacobs, S., et al., 2015. Importance of the GluN2B carboxy-terminal domain for enhancement of social memories. Learn Mem. 22, 401–10.

Jost, B. C., Grossberg, G. T., 1995. The natural history of Alzheimer’s disease: a brain bank study. J Am Geriatr Soc. 43, 1248–55.

Klein, A. S., et al., 2016. Early Cortical Changes in Gamma Oscillations in Alzheimer’s Disease. Front Syst Neurosci. 10, 83.

Kuiper, J. S., et al., 2015. Social relationships and risk of dementia: A systematic review and meta-analysis of longitudinal cohort studies. Ageing Res Rev. 22, 39–57.

Lalonde, R., et al., 2003. Transgenic mice expressing the betaAPP695SWE mutation: effects on exploratory activity, anxiety, and motor coordination. Brain Res. 977, 38–45.

Latif-Hernandez, A., et al., 2017. Subtle behavioral changes and increased prefrontal-hippocampal network synchronicity in APP(NL-G-F) mice before prominent plaque deposition. Behav Brain Res.

Li, X. X., Li, Z., 2018. The impact of anxiety on the progression of mild cognitive impairment to dementia in Chinese and English data bases: a systematic review and meta-analysis. Int J Geriatr Psychiatry. 33, 131–140.

Lin, T. W., et al., 2015. Running exercise delays neurodegeneration in amygdala and hippocampus of Alzheimer’s disease (APP/PS1) transgenic mice. Neurobiol Learn Mem. 118, 189–97.

Liu, Y., et al., 2010. Analysis of regional MRI volumes and thicknesses as predictors of conversion from mild cognitive impairment to Alzheimer’s disease. Neurobiol Aging. 31, 1375–85.

Livingston, G., et al., 2017. Dementia prevention, intervention, and care. Lancet. 390, 2673–2734.

Ly, P. T., et al., 2011. Detection of neuritic plaques in Alzheimer’s disease mouse model. J Vis Exp.

McHugh, S. B., et al., 2004. Amygdala and ventral hippocampus contribute differentially to mechanisms of fear and anxiety. Behav Neurosci. 118, 63–78.

McNally, J. M., et al., 2011. Complex receptor mediation of acute ketamine application on in vitro gamma oscillations in mouse prefrontal cortex: modeling gamma band oscillation abnormalities in schizophrenia. Neuroscience. 199, 51–63.

Middleton, S., et al., 2008. NMDA receptor-dependent switching between different gamma rhythm-generating microcircuits in entorhinal cortex. Proc Natl Acad Sci U S A. 105, 18572–7.

Moy, S. S., et al., 2004. Sociability and preference for social novelty in five inbred strains: an approach to assess autistic-like behavior in mice. Genes Brain Behav. 3, 287–302.

Nadler, J. J., et al., 2004. Automated apparatus for quantitation of social approach behaviors in mice. Genes Brain Behav. 3, 303–14.

Nakazono, T., et al., 2017. Impaired In Vivo Gamma Oscillations in the Medial Entorhinal Cortex of Knock-in Alzheimer Model. Front Syst Neurosci. 11, 48.

Okello, A., et al., 2009. Conversion of amyloid positive and negative MCI to AD over 3 years: an 11C-PIB PET study. Neurology. 73, 754–60.

Okuyama, T., et al., 2016. Ventral CA1 neurons store social memory. Science. 353, 1536–1541.

Palmer, K., et al., 2007. Predictors of progression from mild cognitive impairment to Alzheimer disease. Neurology. 68, 1596–602.

Pervolaraki, E., et al., 2017. Ventricular myocardium development and the role of connexins in the human fetal heart. Sci Rep. 7, 12272.

Pervolaraki, E., et al., 2018. The developmental transcriptome of the human heart. Sci Rep. 8, 15362.

Pervolaraki, E., et al., 2019. The within-subject application of diffusion tensor MRI and CLARITY reveals brain structural changes in Nrxn2 deletion mice. Mol Autism. 10, 8.

Pietrzak, R. H., et al., 2015. Amyloid-beta, anxiety, and cognitive decline in preclinical Alzheimer disease: a multicenter, prospective cohort study. JAMA Psychiatry. 72, 284–91.

Plant, C., et al., 2010. Automated detection of brain atrophy patterns based on MRI for the prediction of Alzheimer’s disease. Neuroimage. 50, 162–74.

Poulin, S. P., et al., 2011. Amygdala atrophy is prominent in early Alzheimer’s disease and relates to symptom severity. Psychiatry Res. 194, 7–13.

Ramakers, I. H., et al., 2013. Anxiety is related to Alzheimer cerebrospinal fluid markers in subjects with mild cognitive impairment. Psychol Med. 43, 911–20.

Saito, T., et al., 2014. Single App knock-in mouse models of Alzheimer’s disease. Nat Neurosci. 17, 661–3.

Sakakibara, Y., et al., 2018. Cognitive and emotional alterations in App knock-in mouse models of Abeta amyloidosis. BMC Neurosci. 19, 46.

Scheff, S. W., Price, D. A., 2001. Alzheimer’s disease-related synapse loss in the cingulate cortex. J Alzheimers Dis. 3, 495–505.

Scheltens, P., et al., 2016. Alzheimer’s disease. Lancet. 388, 505–17.

Schenk, D., et al., 1999. Immunization with amyloid-beta attenuates Alzheimer-disease-like pathology in the PDAPP mouse. Nature. 400, 173–7.

Shin, L. M., Liberzon, I., 2010. The neurocircuitry of fear, stress, and anxiety disorders. Neuropsychopharmacology. 35, 169–91.

Simon, A. M., et al., 2009. Overexpression of wild-type human APP in mice causes cognitive deficits and pathological features unrelated to Abeta levels. Neurobiol Dis. 33, 369–78.

Stujenske, J. M., et al., 2014. Fear and safety engage competing patterns of theta-gamma coupling in the basolateral amygdala. Neuron. 83, 919–33.

Tallon-Baudry, C., et al., 1998. Induced gamma-band activity during the delay of a visual short-term memory task in humans. J Neurosci. 18, 4244–54.

Teri, L., et al., 1999. Anxiety of Alzheimer’s disease: prevalence, and comorbidity. J Gerontol A Biol Sci Med Sci. 54, M348–52.

Tondelli, M., et al., 2012. Structural MRI changes detectable up to ten years before clinical Alzheimer’s disease. Neurobiol Aging. 33, 825 e25–36.

Wang, C. C., et al., 2011. A critical role for GluN2B-containing NMDA receptors in cortical development and function. Neuron. 72, 789–805.

Whyte, L. S., et al., 2018. Reduction in open field activity in the absence of memory deficits in the App(NL-G-F) knock-in mouse model of Alzheimer’s disease. Behav Brain Res. 336, 177–181.

Wilson, R. S., et al., 2007. Loneliness and risk of Alzheimer disease. Arch Gen Psychiatry. 64, 234–40.

