## Supplementary material for "Insoluble Aβ overexpression in an *App* knock-in mouse model alters microstructure and gamma oscillations in the prefrontal cortex, causing anxiety-related behaviours": Supp Materials

**Supplemental Material**

**Supp. Table 1**

| **Brain Region** | **DTI Measure** | **ANOVA Comparison** | **F Value** | **P Value** |
| --- | --- | --- | --- | --- |
| Anterior Amygdala | FA | Genotype | F_(1, 14)_ <1 | P = 0.384 |
|  |  | Hemisphere | F_(1, 14)_ = 2.76 | P = 0.095 |
| Posterior Amygdala | FA | Genotype | F_(1, 14)_ <1 | P = 0.300 |
|  |  | Hemisphere | F_(1, 14)_ <1 | P = 0.184 |
| BLA | FA | Genotype | F_(1, 14)_ <1 | P = 0.268 |
|  |  | Hemisphere | F_(1, 14)_ <1 | P = 0.168 |
| Anterior Amygdala | MD | Genotype | F_(1, 14)_ <1 | P = 0.179 |
|  |  | Hemisphere | F_(1, 14)_ <1 | P = 0.353 |
| Posterior Amygdala | MD | Genotype | F_(1, 14)_ <1 | P = 0.274 |
|  |  | Hemisphere | F_(1, 14)_ <1 | P = 0.226 |
| BLA | MD | Genotype | F_(1, 14)_ = 3.23 | P = 0.084 |
|  |  | Hemisphere | F_(1, 14)_ = 1.55 | P = 0.137 |
| Anterior Amygdala | AD | Genotype | F_(1, 14)_ <1 | P = 0.173 |
|  |  | Hemisphere | F_(1, 14)_ <1 | P = 0.242 |
| Posterior Amygdala | AD | Genotype | F_(1, 14)_ <1 | P = 0.221 |
|  |  | Hemisphere | F_(1, 14)_ <1 | P = 0.252 |
| BLA | AD | Genotype | F_(1, 14)_ = 5.81 | P = 0.053 |
|  |  | Hemisphere | F_(1, 14)_ = 2.55 | P = 0.105 |
| Anterior Amygdala | RD | Genotype | F_(1, 14)_ <1 | P = 0.189 |
|  |  | Hemisphere | F_(1, 14)_ <1 | P = 0.247 |
| Posterior Amygdala | RD | Genotype | F_(1, 14)_ <1 | P = 0.332 |
|  |  | Hemisphere | F_(1, 14)_ <1 | P = 0.194 |
| BLA | RD | Genotype | F_(1, 14)_ = 2.67 | P = 0.100 |
|  |  | Hemisphere | F_(1, 14)_ <1 | P = 0.205 |

Statistical analysis of the anterior (Bregma -1.94 mm) and posterior (Bregma -3.28 mm) amygdala, analysed for fractional anisotropy (FA), mean diffusion (MD), axial diffusion (AD) and radial diffusion (RD). Corrected P values stated (Benjamini-Hochberg corrected).

**Supp. Table 2**

| **Brain Region** | **DTI Measure** | **ANOVA Comparison** | **F Value** | **P Value** |
| --- | --- | --- | --- | --- |
| Anterior Hippocampus | FA | Genotype | F_(1,14)_ <1 | P = 0.342 |
|  |  | Hemisphere | F_(1, 14)_ = 2.18 | P = 0.116 |
| Posterior Hippocampus | FA | Genotype | F_(1, 14)_ <1 | P = 0.263 |
|  |  | Hemisphere | F_(1, 14)_ <1 | P = 0.305 |
| Anterior Hippocampus | MD | Genotype | F_(1, 14)_ = 2.89 | P = 0.089 |
|  |  | Hemisphere | F_(1, 14)_ <1 | P = 0.311 |
| Posterior Hippocampus | MD | Genotype | F_(1, 14)_ = 12.18 | P = 0.011 |
|  |  | Hemisphere | F_(1, 14)_ = 1.07 | P = 0.163 |
| **Anterior Hippocampus** | **AD** | **Genotype** | **F_(1, 14)_ = 7.56** | **P = 0.047** |
|  |  | Hemisphere | F_(1, 14)_ <1 | P = 0.316 |
| Posterior Hippocampus | AD | Genotype | F_(1, 14)_ = 4.45 | P = 0.068 |
|  |  | Hemisphere | F_(1, 14)_ <1 | P = 0.211 |
| Anterior Hippocampus | RD | Genotype | F_(1, 14)_ = 1.52 | P = 0.142 |
|  |  | Hemisphere | F_(1, 14)_ <1 | P = 0.237 |
| **Posterior Hippocampus** | **RD** | **Genotype** | **F_(1, 14)_ <1** | **P = 0.005** |
|  |  | Hemisphere | F_(1, 14)_ <1 | P = 0.158 |

Statistical analysis of the anterior (Bregma -1.94 mm) and posterior (Bregma -3.28 mm) hippocampus for fractional anisotropy (FA), mean diffusion (MD), axial diffusion (AD) and radial diffusion (RD). Corrected P values stated (Benjamini-Hochberg corrected).

**Supp. Figure 1.**

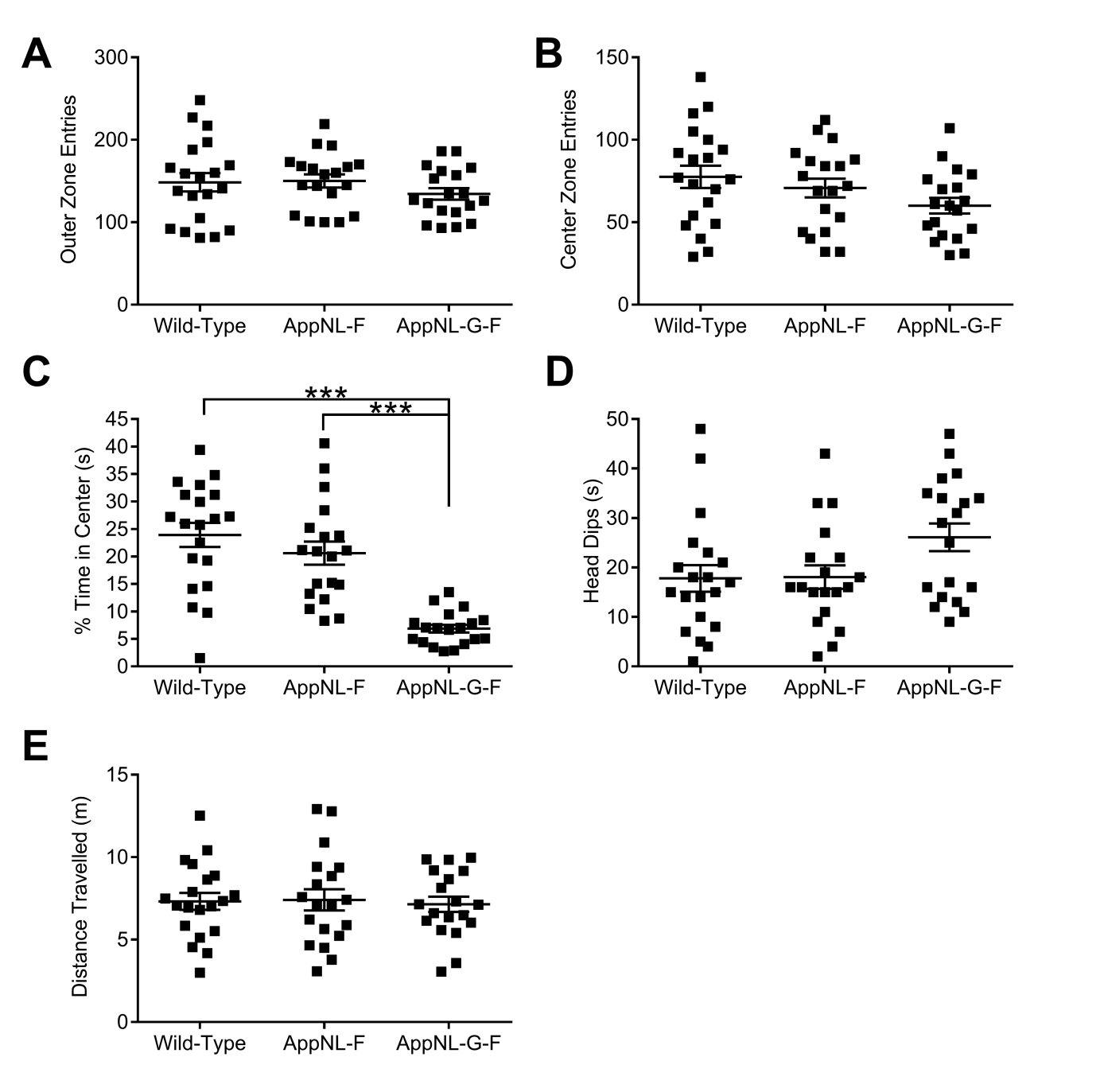

**A.** No significant differences were observed for entries made to the outer zone (genotype F_(2, 55)_ = 1.05, p = 0.357, genotype x sex F_(2, 55)_ <1, p = 0.809). **B.** Similarly, all genotypes made similar entries into the centre zone (genotype F_(2, 55)_ = 2.67, p = 0.079, genotype x sex F_(2, 55)_ <1, p = 0.934). **C.** *App^NL-G-F^* KI mice spent significantly less time in the centre zone in the elevated plus maze (genotype F_(2, 55)_ = 6.14, p = 0.004, sex F_(1, 55)_ <1, p = 0.411, genotype x sex F_(2, 55)_ <1, p = 0.548. Tukey’s post hoc: wild-type vs. *App^NL-G-F^* KI mice p = 0.061, *App^NL-F^* KI mice vs. *App^NL-G-F^* KI mice p = 0.004). **D.** Although there was a main effect of genotype, the number of head dips did not differ between the genotypes in the elevated plus maze (genotype F_(2, 58)_ = 3.76, p = 0.03, sex F_(1, 55)_ = 2.67, p = 0.108, genotype x sex F_(2, 55)_ <1, p = 0.843. Tukey’s post hoc: wild-type vs. *App^NL-G-F^* KI mice p = 0.054, *App^NL-F^* KI mice vs. *App^NL-G-F^* KI mice p = 0.068). **E.** Distance travelled in the elevated plus maze was similar between the genotypes (genotype F_(2, 55)_ <1, p = 0.953, sex F_(1, 55)_ <1, p = 0.971, genotype x sex F_(2, 55)_ <1, p = 0. 396).

**Supp. Figure 2**

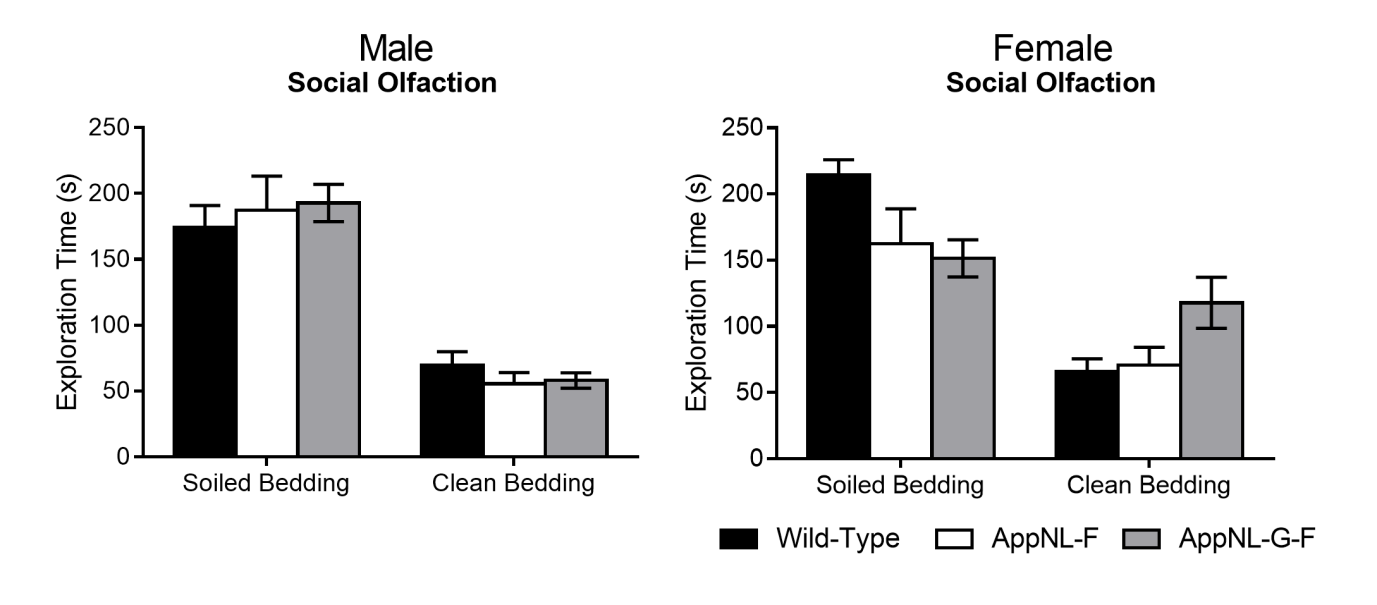

Exploration of cages containing social smells (soiled bedding) and non-social smells (clean bedding). Although all genotypes showed preference for exploring the social smell, there was a discrimination difference depending upon genotype and sex (see main text for statistics).

**Supp. Figure 3**

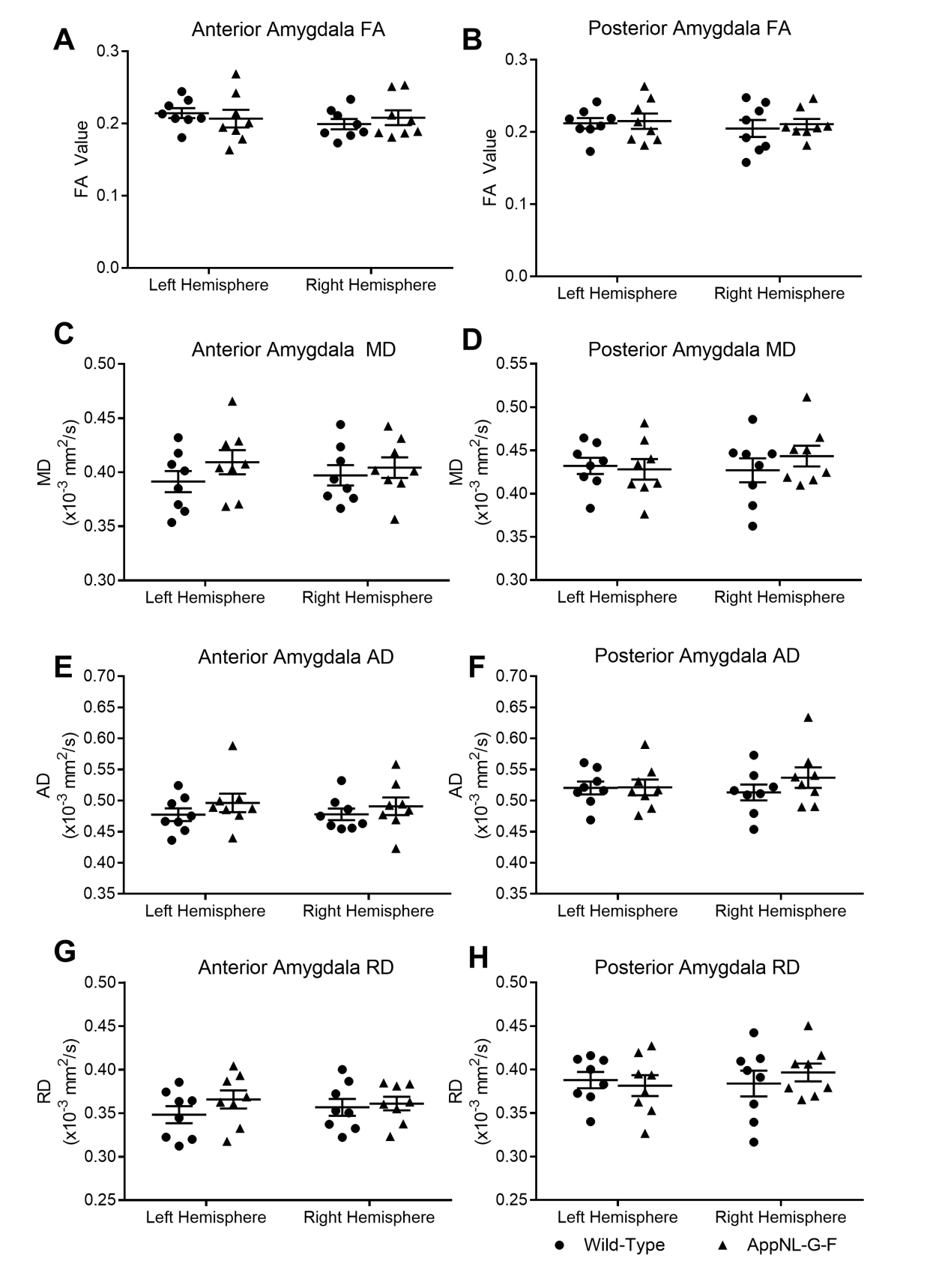

Fractional anisotropy (FA), mean diffusivity (MD), axial diffusivity (AD) and radial diffusivity (RD) in the amygdala. The entire amygdala was segmented from DTI images at two regions; anterior (**A,** **C, E, G**: Bregma -1.94 mm) and posterior (**B, D, F, H**: Bregma -2.46 mm). No significant differences were observed between the genotypes for anterior or posterior regions. Error bars represent s.e.m. Wild-type n=8, *App^NL-G-F^* KI n=8.

**Supp. Figure 4**

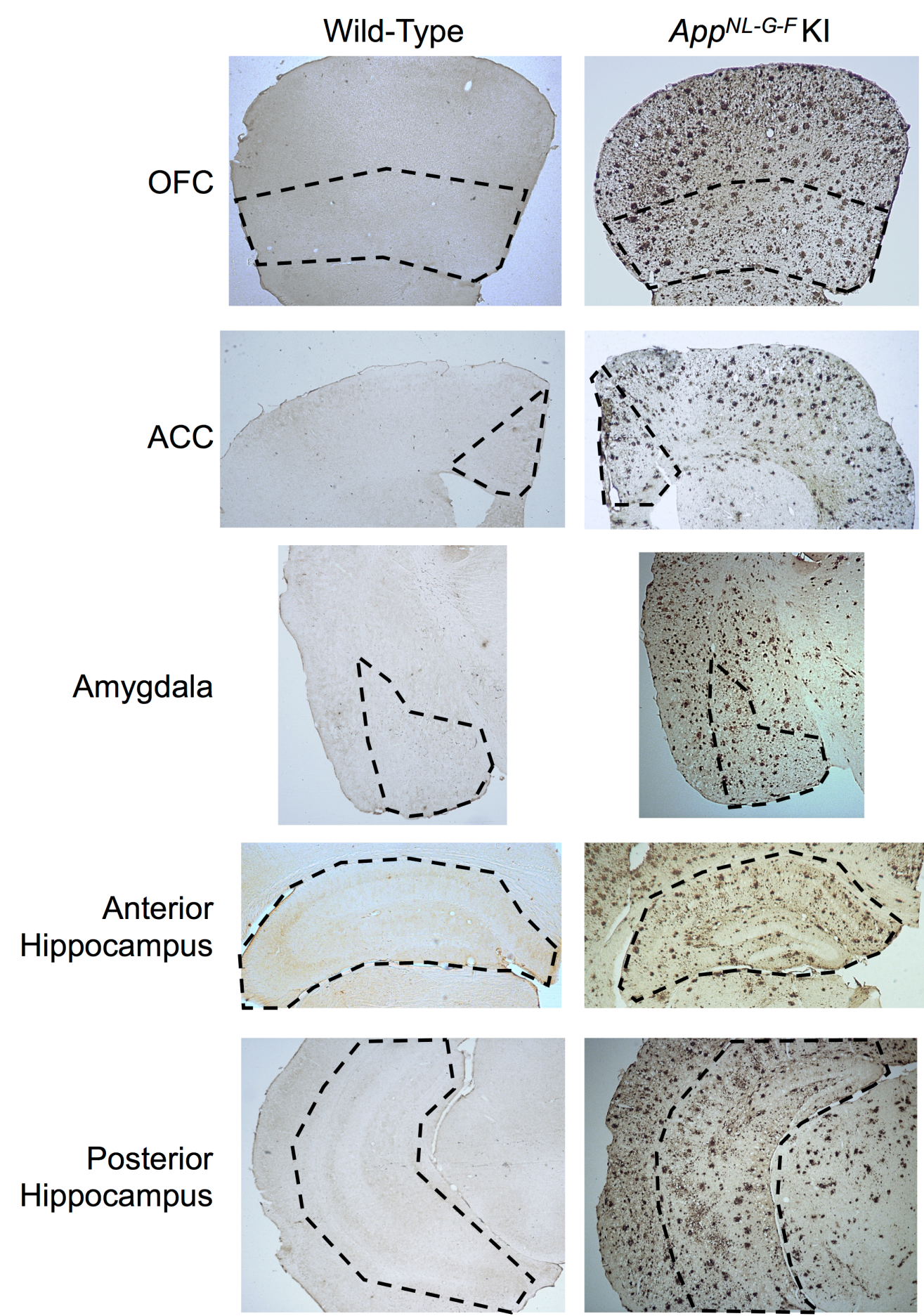

Amyloid plaque staining in wild-type and *App^NL-G-F^* KI mice in the orbitofrontal cortex (OFC), the anterior cingulate cortex (ACC) the amygdala, and the anterior and posterior hippocampus. Dashed line represents the brain regions of interest.
